# Human-specific CCG-repeat insertion in the *RBFOX1* promoter implicated in autism and speech enhances *EGR1*-responsive activity

**DOI:** 10.64898/2026.04.21.719679

**Authors:** Chul Lee, Matthew H. Davenport, Erich D. Jarvis

**Author notes:** Correspondence (E.D.J.). These authors contributed equally (Chul Lee, Matthew H. Davenport).

## Abstract

Human-specific promoter innovations have been predicted to shape human-specific traits. Here, we investigated the *RBFOX1* splice regulator gene, which is specifically upregulated in human and songbird speech/song brain areas. We found that the human *RBFOX1* promoter associated with autism and language impairment harbors a human-specific CCG trinucleotide insertion creating a functional CCG-repeat motif. Comparative single-nucleus transcriptome analyses of primate brains indicate that the human promoter drives differential expression in cortical excitatory neurons. Reporter assays show that the human CCG-repeat motif, relative to the chimpanzee allele, increases promoter activity with the *EGR1* transcription factor. Genome-wide, the CCG-repeat motif marks core promoters of 104 genes, related to postsynaptic function and autism. In *PTCHD1*, a clinical duplication in autism patients creates another CCG-repeat motif that similarly increases *EGR1*-responsive promoter activity. Our findings suggest that small CCG-repeat variants in regulatory regions can exert large effects on gene regulation and apparent associated traits.

## Introduction

Understanding the causal variants and molecular mechanisms underlying human-specific traits and diseases is still a central challenge in the new era of near-complete genome sequences^1–3^. In the case of language, this trait is associated with specialized up-or down-regulation of hundreds of genes within speech-producing brain regions relative to their non-vocal surrounding tissue and to analogous convergent brain regions in song-learning birds^4,5^. Among these genes, the RNA-binding protein Fox1 homolog (*RBFOX1*; also known as *A2BP1* and *FOX1*) showed the most significantly differential expressions in glutamatergic neurons of the songbird brain^6^. *RBFOX1* regulates cell-type-specific alternative splicing, synaptic gene expression programs, and neuronal excitability^7–11^.

*RBFOX1*-disrupting variants and copy-number changes have been reported in individuals with ASD, intellectual disability, epilepsy, and related neurodevelopmental or psychiatric phenotypes^12–16^. Further, ASD patients’ brains show reduced *RBFOX1* expression and dysregulation of *RBFOX1*-dependent alternative splicing^17^. *RBFOX1* also uses multiple tissue-specific alternative promoters to generate distinct transcript isoforms^18,19^. A previously reported individual with ASD and language impairment carried a large deletion encompassing an alternative *RBFOX1* promoter and showed selective reduction of the corresponding transcript isoform^12,13^. The deletion established the functional relevance of the promoter-containing interval but did not identify the sequence feature responsible for its activity or whether that feature was human-derived. Therefore, we treated this speech-related and ASD-risk promoter as a starting point for identifying human-specific functional elements.

However, identifying human-derived sequence features within this *RBFOX1* promoter poses two methodological challenges. First, comparative approaches for human regulatory innovation prioritize alignable sequences with accelerated substitution rates^20–22^, underrepresenting short lineage-specific insertions or repeat-copy changes. For *RBFOX1*, such approaches have identified human-accelerated enhancers elsewhere in the locus^23^, but not in the ASD-risk promoter. Second, repeat-rich promoter sequences were also frequently collapsed, misassembled, erroneously duplicated, or absent in legacy short-read genome assemblies^24–27^. However, high-quality long-read genome assemblies of the Vertebrate Genomes Project (VGP)^27,28^ and associated consortia^29–31^ now enable base-resolved comparisons of these regions across species and populations, complemented by genomic evidence from archaic hominins^32^.

Here, we tested this hypothesis by integrating long-read-assembled genomes from the VGP Phase 1 project, comparative genomics approaches, human variant-effect estimates, and promoter reporter assays. In the *RBFOX1* promoter, we identified a human-lineage-specific CCG trinucleotide insertion that created a CCG-repeat-binding motif for the *EGR1* transcription factor, thereby increasing brain *RBFOX1* expression in *in silico* variant-effect estimates using AlphaGenome^61^ and enhancing promoter activity in *in vitro* reporter assays. Moreover, in a promoter of the *PTCHD1* gene, another gene with an ASD-associated duplication^33^ resulting in another CCG repeat motif, also enhanced *EGR1* responsiveness. These findings suggest that CCG-repeated sequences serve as promoter elements linking human-specific regulatory evolution to susceptibility to neurodevelopmental disease.

## Results

### A disease-susceptible promoter of *RBFOX* with a human-specific CCG insertion

A previously reported individual with autism and language impairment carries a 208-kb deletion (chr16:6,002,834-6,210,814 in hg38, liftover from hg19 to hg38) that removed an alternative *RBFOX1* promoter and selectively reduced the corresponding transcript isoform^12,13^ (**Figure 1A**; **Table S1**). Focusing on the deleted promoter, we searched clinical structural variants associated with neurodevelopmental deficits related to autism and language dysfunction using the DECIPHER database^34^ and found 17 variants reported in 36 individuals (**Figure 1A**; **Table S1**). Based on transcription start site (TSS) peaks from FANTOM5^35–37^ and candidate cis-regulatory element (cCRE) annotations from ENCODE4^38^, we marked an active initiation region within this promoter (**Figure 1B**). Additionally, we analyzed comparative single-cell transcriptome patterns of the primary motor cortex of human, macaque, and marmoset^39^ and observed human-increased promoter-proximal coverage in layer 5 (L5) and L6 excitatory neurons, which was weak or absent in homologous non-human primate neurons (**Figure 1B**).

**Figure 1.**
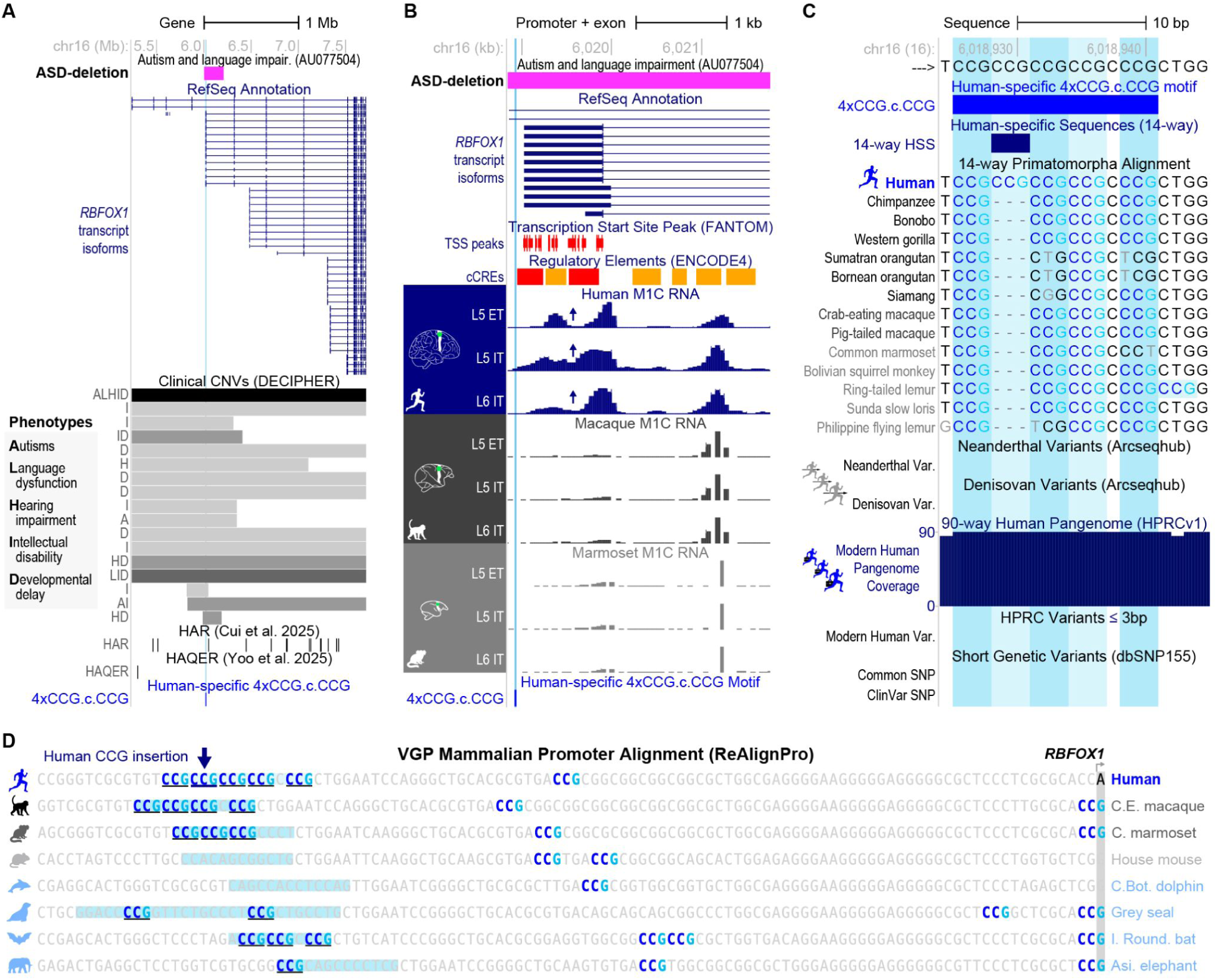
Clinical structural variants include an *RBFOX1* promoter-containing human-lineage CCG insertion. (**A**) Gene-wide UCSC view of *RBFOX1* showing RefSeq isoforms, the autism-associated AU077504 deletion, DECIPHER CNVs, HAR/HAQER annotations, which do not overlap the motif, and the human-specific 4xCCG.c.CCG motif at chr16:6,018,926-6,018,941 (hg38, https://genome.ucsc.edu/s/clee03/CCG2026_Fig1A_ASDdel). The darkness level of gray bars represents the density of the number of clinical phenotypes associated with each structural variant. Phenotype codes: L, language dysfunction; H, hearing impairment; A, autism; I, intellectual disability; D, developmental delay. Detailed information on clinical variants is in **Table S1**. (**B**) Promoter-level UCSC view supported by FANTOM5 CAGE peaks and ENCODE cCREs, with primary motor cortex single-nucleus RNA-seq coverage across excitatory neuron subtypes in human, macaque, and marmoset. Arrows indicate human-enriched promoter-proximal transcript heterogeneity (https://genome.ucsc.edu/s/clee03/CCG2026_Fig1B_Exp). (**C**) Sequence-level UCSC view with the 14-way primatomorpha alignment showing the unique CCG insertion in the human genome (https://genome.ucsc.edu/s/clee03/CCG2026_Fig1C_Seq). **Figure S1A** summarizes other publicly accessible alignments. Neanderthal, Denisovan, HPRC pangenome, dbSNP155, and ClinVar tracks indicate conservation between archaic hominins and modern humans and no common or clinical SNPs overlapping the motif. (**D**) ReAlignPro mammalian promoter alignment anchored at the human *RBFOX1* TSS. Conserved flanking bases, CCG-repeat bases, and the homologous regions to the human 4xCCG.c.CCG motifs are indicated in a more visible colored font. **Figure S1B** summarizes the raw alignment with gaps as sequence logos, and **Table S2** shows the TSS-anchored gapless alignment of 145 VGP assemblies. Orthologous sequences of the human 4xCCG.c.CCG motif were marked with sky-blue backgrounds. The human-specific CCG insertion was marked using a navy arrow. The workflow of ReAlignPro is shown in **Figure S2**. See also **Figures S1** and **S2**, and **Tables S1** and **S2**.

To test whether the promoter contained any human-derived genomic elements, we constructed a 14-way primatomorpha genome-wide alignment using long-read-based, high-quality assemblies of 13 primates and 1 flying lemur outgroup (Dermoptera) from the VGP^40^ and Telomere-to-Telomere (T2T) consortiums^31^ (**STAR Methods**). In this alignment, we resolved a human-specific CCG trinucleotide repeat at chr16:6,018,926–6,018,928 (hg38), represented by three nucleotide gaps at the homologous position in all 13 non-human animal sequences (**Figure 1C**). The insertion completes a five-copy CCG-repeat motif (CCGCCGCCGCCGcCCG: four CCG units, an interrupting cytosine ‘c’, and a fifth CCG; hereafter the expanded human allele, 4xCCG.c.CCG). Except for the insertion, the human promoter sequence is otherwise identical to that of the closest relatives, chimpanzee, bonobo, and gorilla (**Figure 1C**). The ring-tailed lemur is the only other sampled species with five copies, but it had an interrupting ‘c’ after the 3rd CCG repeat (CCGCCGCCGcCCGCCG, 3xCCG.c.2xCCG; **Figure 1C**).

Because acceleration screens score substitution rates within alignable windows, a short lineage-specific insertion of this kind falls outside their detection model, and neither Human Accelerated Region (HAR)^20,22^ nor Human Ancestor Quickly Evolved Region (HAQER)^21,22^ catalogs annotate this promoter (**Figure 1A**). Our analyses of independent T2T ape, CACTUS, and MULTIZ genome-wide alignments in the UCSC genome browser associated with the T2T and VGP projects^31,41^ consistently support an expanded human configuration despite alternative gap placement within the repeat (**Figure S1A**). To resolve the technical alternatives for repetitive regions by anchoring promoter sequences at the TSS, we developed ReAlignPro (**STAR Methods**, **Figure S2**). We queried 153 high-quality VGP non-human mammalian genome assemblies^40^ with the human promoter sequence and recovered *RBFOX1* orthologous promoters from 143 of them (**Table S2**). It placed the human CCG insertion, which was absent in all non-human mammals surveyed, including non-human vocal-learning clades (**Figure 1D**; **Table S2**).

In archaic hominins, genomic sequences in the two Arcseqhub datasets^42^ and the four Allen Ancient DNA Resource (AADR) v66.p1 datasets^32^ contained the same *RBFOX1* CCG repeat motif as modern humans, with no nucleotide polymorphism (SNPs; **Figures 1C** and **S3A**). We also examined published BAM alignments of high-coverage ancient hominin genomes of three Neanderthals (Altai, Vindija 33.19, and Chagyrskaya 8) and of one Denisovan (Denisova 3). After quality filtering and requiring exact 10-bp anchors on both sides of the motif, 16 and 2 genomic reads supported the exact human-reference CCG repeat sequence, respectively. Only Vindija contained three reads with C-to-T mismatches in the CCG repeat, consistent with post-mortem deamination^43^ (**Figure S3B-D**). In modern humans, the CCG motif was invariant across the 90 high-quality haplotypes sequenced by the Human Pangenome Reference Consortium (HPRC) assemblies^30^, with no common dbSNP155^44^ variants or ClinVar variants^45^ within it (**Figure 1C**).

The primate alignment and the archaic hominin reads together bracket the origin of the allele: the cross-species alignment places its appearance after the Chimpanzee-Human divergence (6.4 Mya)^46^, and the archaic read-level evidence places it before the divergence of modern from archaic hominins.

### The human insertion creates a functional motif for *EGR1*-responsive activity

Scanning the human-specific CCG-repeat region with 3 bp flanking sequences (tgtCCGCCGCCGCCGcCCGctg) in the human genome with JASPAR CORE 2022 position-weight models^47^, we predicted two binding sites for the *EGR1* transcription factor and one for the *WT1* transcription factor, which is a paralogous gene of *EGR1*^48–50^ (**Figure 2A; Tables S2 and S3)**. *EGR1* is a well-known activity-dependent transcription factor in the human brain with a C2H2 zinc-finger binding domain and is upregulated in specific neuron types when they exhibit increased synaptic activity^51–54^. Its paralog *WT1* has DNA-binding properties that partially overlap with those of *EGR1*^55^. The human CCG insertion created a 14-bp predicted *EGR1* binding site (CGCCGCCGCCGcCC), present only in the human allele among primates, and is 3 bp upstream of another 14-bp predicted *EGR1* and *WT1* binding site (gtCCGCCGCCGCCG), which differs from the non-human primate version only by the last base pair (gtCCGCCGCCGcCC). The ring-tailed lemur allele (tgtCCGCCGCCGcCCGCCG), despite carrying five CCG copies, has an interrupting ‘c’ that leaves an uninterrupted run of only three CCG units and, like the four-copy non-human great-ape alleles, yields a single predicted *EGR1* site (gtCCGCCGCCGcCC) identical to the chimpanzee version and no *WT1* site (**Table S3**). There were two other 3’ locations in the promoter with predicted *EGR1* and *WT1* binding sites, but none of these were specific to humans (**Figure 2A**).

**Figure 2.**
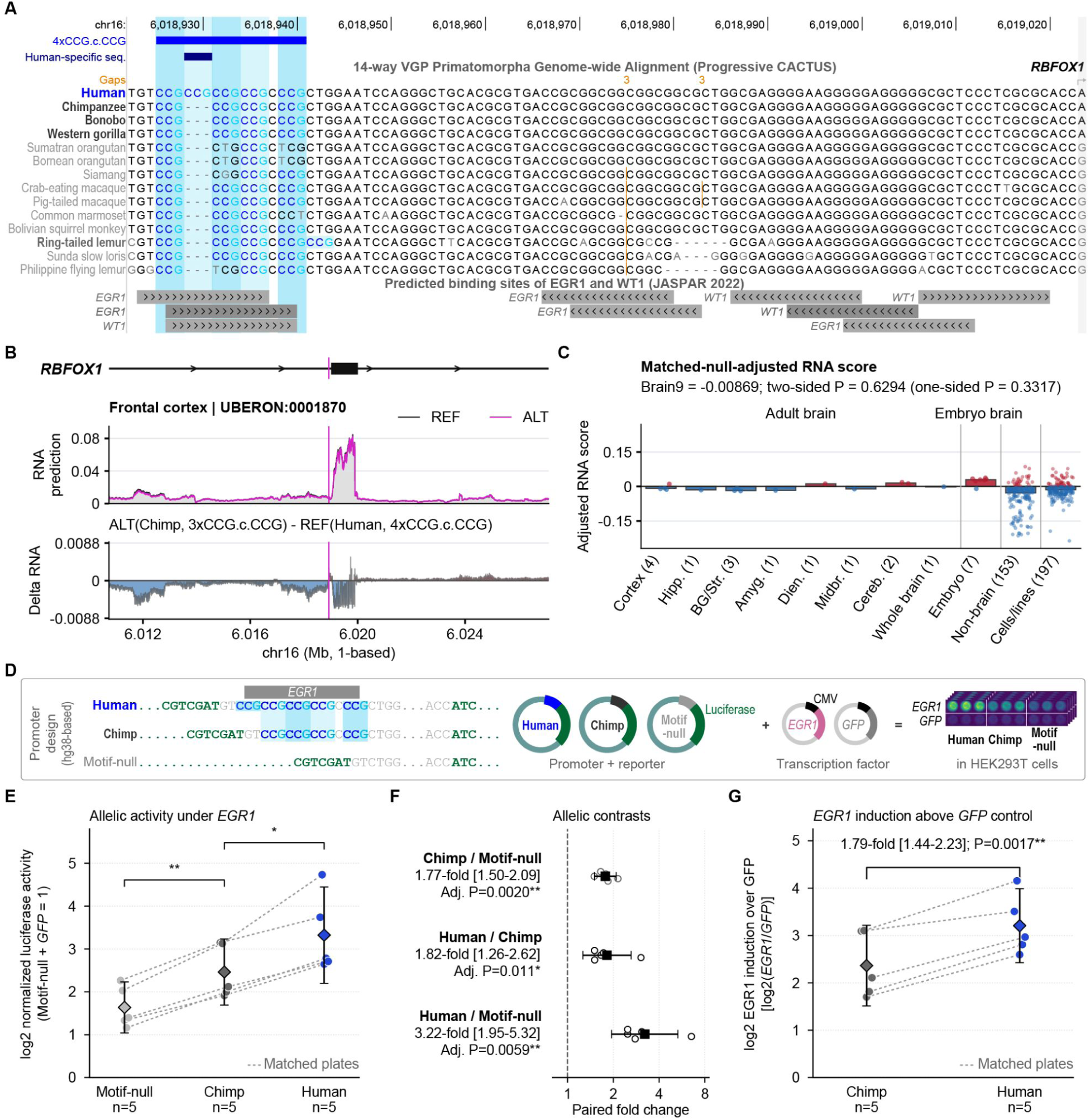
The human *RBFOX1* CCG-repeat allele increases *EGR1*-responsive promoter activity. (**A**) 14-way VGP Primatomorpha alignment and JASPAR CORE 2022 motif scan of the alternative *RBFOX1* core promoter (chr16:6,018,923-6,019,024, + strand, https://genome.ucsc.edu/s/clee03/CCG2026_Fig2A_PromoterSeq_JASPAR). Within the human CCG-repeat allele with 3-bp up/down stream flanking sequences (tgtCCGCCGCCGCCGcCCGctg), the human-specific CCG insertion generates one predicted *EGR1* binding site (CGCCGCCGCCGcCC), which is uniquely observed in humans, and one predicted *EGR1* and *WT1* binding site (gtCCGCCGCCGCCG), which is almost identical to the chimpanzee allele (gtCCGCCGCCGCCC; **Table S3**). (**B**) AlphaGenome-predicted whole-brain RNA-seq profiles for the human hg38 reference allele (REF) and the chimpanzee-like 3-bp deletion allele (ALT) in the human genomic context. ΔRNA is defined as ALT minus REF. (**C**) Matched-null-calibrated AlphaGenome RNA effects summarized by tissue or cell context using Alpha-GENIE, which was extended for this study to support matched pseudo-deletion calibration and context-based track aggregation. Negative values indicate lower predicted RNA output from ALT; numbers in parentheses indicate the number of tissue/cell types. Calibration and grouping procedures are described in **STAR Methods**. (**D**) Luciferase reporters containing the human, chimpanzee-like, or motif-null promoter alleles, co-transfected into HEK293T cells with CMV-*EGR1* or a CMV-*GFP* expression-vector control. Additional reporter and transcription-factor controls are shown in **Figure S4A**. (**E**) Plate-matched promoter activity under *EGR1*. The luciferase signal was background-subtracted, normalized within the plate to motif-null+*GFP* levels, averaged across technical replicates, and log2-transformed. Points represent five independent transfections; dashed lines connect alleles assayed on the same plate, and diamonds with bars show mean log2 activity and 95% confidence intervals. Brackets indicate the two sequential adjacent allele comparisons; significance was determined using the same Holm-adjusted P values reported in (**F**). Additional controls are shown in **Figures S4A,B**. (**F**) Paired allelic fold changes from (**E**). Open circles show plate-level ratios; filled squares and horizontal bars show geometric mean fold changes and 95% confidence intervals. The p-values were obtained from two-sided paired t-tests on log2 ratios and adjusted using the Holm method within the pre-specified three-contrast *EGR1* family. We used the Holm method over the classical Bonferroni method, because it offers more statistical power in correcting the family-wise error rate. (**G**) *EGR1* induction relative to the *GFP*-vector control, calculated for each allele as log_2_(*EGR1*/*GFP*). The allele-by-*EGR1* interaction is the difference between human and chimpanzee induction values, equivalent after back-transformation to (Human*_EGR1_*/Human*_GFP_*)/(Chimp*_EGR1_*/Chimp*_GFP_*). The interaction p-value was obtained from a separate two-sided paired t-test and was not included in the Holm family adjustment in (**F**). See also **Figure S4** and **Tables S2, S3**.

To estimate the effect of the CCG insertion in humans, we used the human hg38 genome assembly allele as the reference (REF) and the chimpanzee-like allele as the alternate (ALT), and computed matched-null-calibrated AlphaGenome RNA effects across tissues using Alpha-GENIE (https://alpha-genie.net; calibration and aggregation in **STAR Methods**). To reduce generic indel-shift effects, we subtracted the median effect of matched 3-bp pseudo-deletions scored in the same genomic context as the null effect model. The calibrated group-level effect (ALT - REF) was negative for the chimpanzee allele in six adult brain-tissue categories except for diencephalon and cerebellum, whereas embryonic brain, non-brain, and cell culture-based groups were heterogeneous (**Figure 2B,C**). That is, the chimpanzee-like contraction was predicted to reduce *RBFOX1* RNA expression across six adult brain-tissue categories in the human genomic context.

To test whether the *RBFOX1* promoter alleles differ in *EGR1* responsiveness, we performed luciferase reporter assays using plasmid constructs carrying the human five-copy, the chimpanzee four-copy, or an engineered null promoter motif, co-transfected into human HEK293T cells with CMV-*EGR1* or CMV-*GFP* expression vectors (**Figure 2D**; **Figure S4A**). As predicted, promoter activity followed this order: motif-null < chimpanzee < human under *EGR1* co-transfection conditions (**Figure 2E**). The chimpanzee reporter was 1.77-fold more active than motif-null (95% CI, 1.50-2.09; Holm-adjusted P = 0.002), the human reporter was 1.82-fold more active than chimpanzee (95% CI, 1.26-2.62; Holm-adjusted P = 0.011), and 3.22-fold more active than motif-null (95% CI, 1.95-5.32; Holm-adjusted P = 0.0059; **Figure 2F**).

To determine whether the human *RBFOX1* promoter allele showed greater *EGR1* responsiveness than the chimpanzee-like allele, we compared *EGR1*-induced fold activation between the two alleles. For each allele, we divided reporter activity under *EGR1* co-transfection by activity under the matched *GFP* control, using that allele’s own control activity as its baseline. This fold activation was 1.79 times higher for the human allele than for the chimpanzee-like allele (95% CI, 1.44-2.23; paired P = 0.0017; **Figure 2G**). Thus, the human promoter allele responded more strongly to *EGR1*.

In additional control tests, *EGR1* lacking its zinc-finger DNA-binding domain (*EGR1ΔBD*) abolished the hierarchy (**Figures S4A, B; Table S3**). *WT1* co-transfection produced a smaller separation in the same order and was similarly affected by the deletion of the DNA-binding domain. The lemur *RBFOX1* allele did not drive human-like expression, consistent with its single predicted *EGR1* site, like the chimpanzee *RBFOX1* allele.

These findings confirmed that the human-specific CCG insertion in the *RBFOX1* promoter created an additional *EGR1*-responsive element, increasing promoter sensitivity.

### The *RBFOX1* CCG-motif recurs at neural and ASD-associated promoters

We next asked how often this CCG repeat configuration occurs elsewhere in the genome, and what other candidate genes it regulates similarly. A BLAST search^56^ identified 266 loci in the hg38 human genome carrying the exact human *RBFOX1* allele sequence (5’-CCGCCGCCGCCGcCCG-3’). Of these, 104 overlapped with core promoters (TSS ± 150 bp 5’ and 3’) of other genes, with four occurring in multi-gene promoter regions (**Figure 3A**; **Table S4**). Motif-bearing promoter loci were distributed across all autosomes and the X chromosome, including multiple loci mapping near *RGPD1* and *RGPD2*, members of a segmentally duplicated gene family on chromosome 2^57^. Functional enrichment analysis found that these genes were enriched for neurodevelopmental and synaptic gene function, including nervous system development, neurogenesis, neuronal differentiation, postsynapse, dendrite, and postsynaptic density terms (**Figure 3B**; **Table S4**).

**Figure 3.**
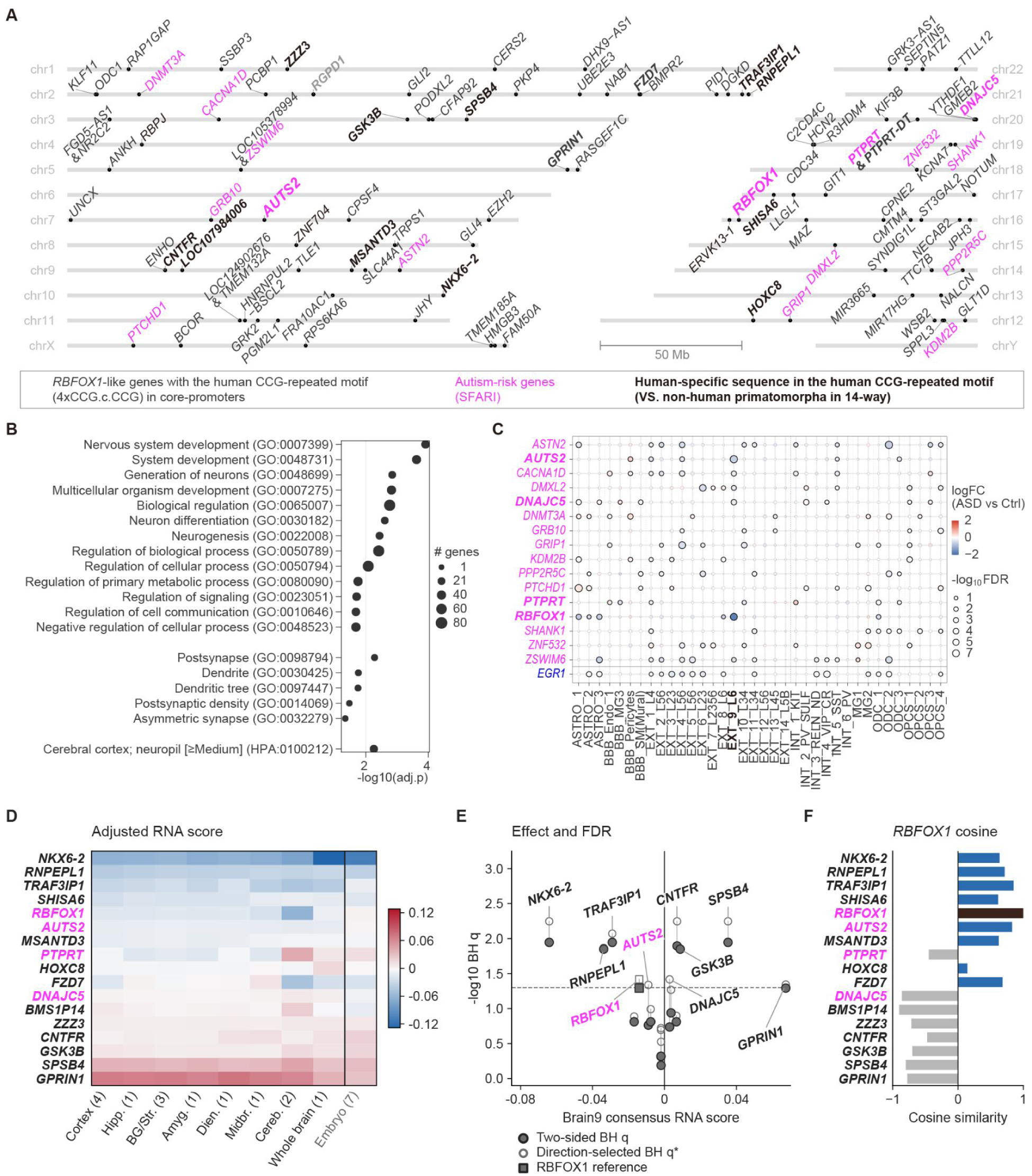
CCG-repeat promoter architecture extends to other neural and ASD-associated genes. (**A**) Chromosomal distribution of human core promoters, defined as TSS ±150 bp for each transcript isoform, carrying the *RBFOX1* human CCG-repeat motif (4xCCG.c.CCG) in hg38 (see also in **Table S4**). Gene labels indicate motif-bearing loci; magenta labels mark SFARI ASD-risk genes, and bold labels mark genes with human-specific sequences on their human CCG-repeat motif. (**B**) Enrichment analysis of motif-bearing genes. Selected GO biological processes, GO cellular components, and Human Protein Atlas terms are shown. Dot position indicates the-log10 adjusted *p-*value, and dot size indicates the number of annotated genes. (**C**) PsychENCODE frontal-cortex single-nucleus RNA-seq differential expression^59^ for ASD-implicated (4xCCG.c.CCG) motif-bearing genes. Rows indicate genes, and columns indicate major cell-type clusters. Dot color shows log2 fold change in ASD versus control, dot size shows-log10 FDR, and outlined dots denote FDR < 0.05. (**D**) AlphaGenome-predicted RNA effects for 20 human-derived CCG-repeat promoter candidates, shown as adjusted ΔRNA for the chimp-like allele relative to the human allele across brain regions, cell types, or cell lines. For deletion alleles, adjusted values incorporate matched-null indel calibration. (**E**) Consensus brain-adjusted ΔRNA versus-log10(adjusted value) for the 17 candidates, showing two-sided BH-adjusted P values and exploratory BH-adjusted observed-direction tail values. (**F**) Cosine similarity of candidate brain RNA-Seq-effect profiles to *RBFOX1*, computed across brain regions, cell types, or cell lines. Panels D-F were generated and modified from Alpha-GENIE (https://alpha-genie.net#multi). See also **Table S4**.

To assess the relevance of neurodevelopmental disease more directly, we intersected the 104 motif-bearing genes with ASD-risk genes in the Simons Foundation Autism Research Initiative (SFARI) database^58^ and identified 16 overlapping genes inclusive of *RBFOX1*: *ASTN2, AUTS2, CACNA1D, DMXL2, DNAJC5, DNMT3A, GRB10, GRIP1, KDM2B, PPP2R5C, PTCHD1, PTPRT, RBFOX1, SHANK1, ZNF532,* and *ZSWIM6*. In our reanalysis of the PsychENCODE frontal-cortex single-nucleus RNA-seq dataset in the ASD scGENE Portal^59^, all 16 motif-bearing SFARI genes were differentially expressed in ASD relative to control individuals in at least one of 35 cell-type clusters (**Figure 3C**). Two 5xCCG motif-bearing SFARI genes with the strongest cell-type-restricted signal were *RBFOX1* and *AUTS2*, and both were downregulated in the same layer 6 excitatory-neuron cluster (EXT_9_L6); for *RBFOX1,* this was the largest effect in the gene set and matched, in direction, the reduced alternative-isoform expression in the deletion ASD carrier.

### Human-derived motif promoters and a convergent candidate module

A sequence match identifies where the CCG-repeat motif occurs but not when it arose. We therefore projected all 266 human loci onto the 14-way VGP primatomorpha alignment and found that 62 contained at least one human-specific base, and 20 of these fell within core promoters (**Figure 3A**, bold; **Table S4**). Three *RGPD*-associated loci lacked a confidently aligned chimpanzee sequence. We interpreted these cases cautiously because recurrent duplication and gene conversion within the *RGPD* gene family complicate one-to-one paralog assignment across ape genomes^60^, so we excluded these three ambiguous *RGPD* loci from the following analysis. The extent of human-specific sequence among the remaining 17 loci varied from 1 to 16 bp.

For each of the 17 loci above, we computed matched-null-calibrated variant effects using AlphaGenome^61^ for the human allele (REF) versus the chimpanzee-like allele (ALT) in human brain tissues. One group of six genes (*NKX6-2, RNPEPL1, TRAF3IP1, SHISA6, RBFOX1, AUTS2,* and *MSANTD3*) showed predicted-upregulation by the human-specific promoter alleles relative to chimpanzee alleles in all 9 categories of brain regions, while another five genes (ZZZ3, *CNTFR, GSK3B, SPSB4, GPRIN1*) showed predicted-downregulation (**Figure 3D**; adjusted ΔRNA ALT - REF, negative = lower predicted output from the chimpanzee allele). Statistical analyses found that *NKX6-2, RNPEPL1,* and *TRAF3IP1* reached FDR_two-sided_ < 0.05 with upregulation in the human brain, and *CNTFR*, *GSK3B,* and *SPSB4* reached FDR_two-sided_ < 0.05 with downregulation in the human brain relative to the chimpanzee (**Figure 3E**). *RBFOX1* and *AUTS2* met the one-sided FDR threshold but not the two-sided one. Lastly, to identify loci with predicted allelic-effect patterns resembling those of *RBFOX1*, we calculated cosine similarity between the matched-null-calibrated RNA-effect profile (ALT - REF) of each locus and that of *RBFOX1* across 9 brain tissue categories of adults and embryos. Eight of the 16 loci showed positive similarity, including seven with scores > 0.5: *TRAF3IP1, AUTS2, RNPEPL1, FZD7, NKX6-2, MSANTD3*, and *SHISA6* (**Figure 3F**). We then prioritized loci using three criteria applied uniformly to all 17 loci: a negative consensus brain RNA effect (ALT - REF) with FDR_one-sided_ < 0.05, positive cosine similarity to *RBFOX1* (> 0), and SFARI annotation^58^. *AUTS2* was the only locus, besides *RBFOX1* itself, that satisfied all three criteria. Further, in the PsychENCODE frontal-cortex single-nucleus RNA-seq dataset^59^, independent of variant-effect predictions, *AUTS2* and *RBFOX1* were concomitantly downregulated in the same layer 6 neuron cluster in autism patients compared to healthy controls (**Figure 3C**).

This screen prioritized candidate genes with promoters similar to *RBFOX1* and predicted the direction and magnitude of gene expression effects based on the entire promoter sequence rather than on the CCG repeat itself. It also does not demonstrate co-regulation. We therefore tested potential promoter activity for co-regulation.

### An ASD-associated *PTCHD1* promoter duplication adds another 5xCCG motif and increases *EGR1*-driven responsive promoter activity

Because the genome-wide analysis indicated that CCG-repeat effects depend on local promoter context, we sought an independent patient-associated repeat gain to test experimentally. *PTCHD1*, an X-linked autism and intellectual-disability risk gene^62–66^, met these criteria: it carries the same 4xCCG.c.CCG motif in its core promoter (**Figure 3A**; **Table S4**) and among the 104 motif-bearing core promoters, is the promoter at which a disease-associated repeat-copy gain – rather than a deletion – has been reported^33^.

ReAlignPro resolved a broadly conserved CCG-rich architecture at the *PTCHD1* core promoter (chrX:23,334,294-23,335,303; **Figure 4A**; **Table S2**). Unlike in *RBFOX1*, the 4xCCG.c.CCG motif in *PTCHD1* is commonly shared in 117 of 151 placental mammal genomes (77.5%). A recurrent 27-bp duplication, reported in three unrelated individuals with ASD and absent from 590 controls^33^, adds an uninterrupted 5xCCG run (CCGCCGCCGCCGCCG) approximately 30 bp upstream of the shared motif (**Figure 4A**). This duplication has been assayed once before, in an extended upstream construct without a supplied transcription factor, in which it reduced basal reporter activity^33^. The original paper did not consider whether it alters responsiveness to *EGR1* at the core promoter.

**Figure 4.**
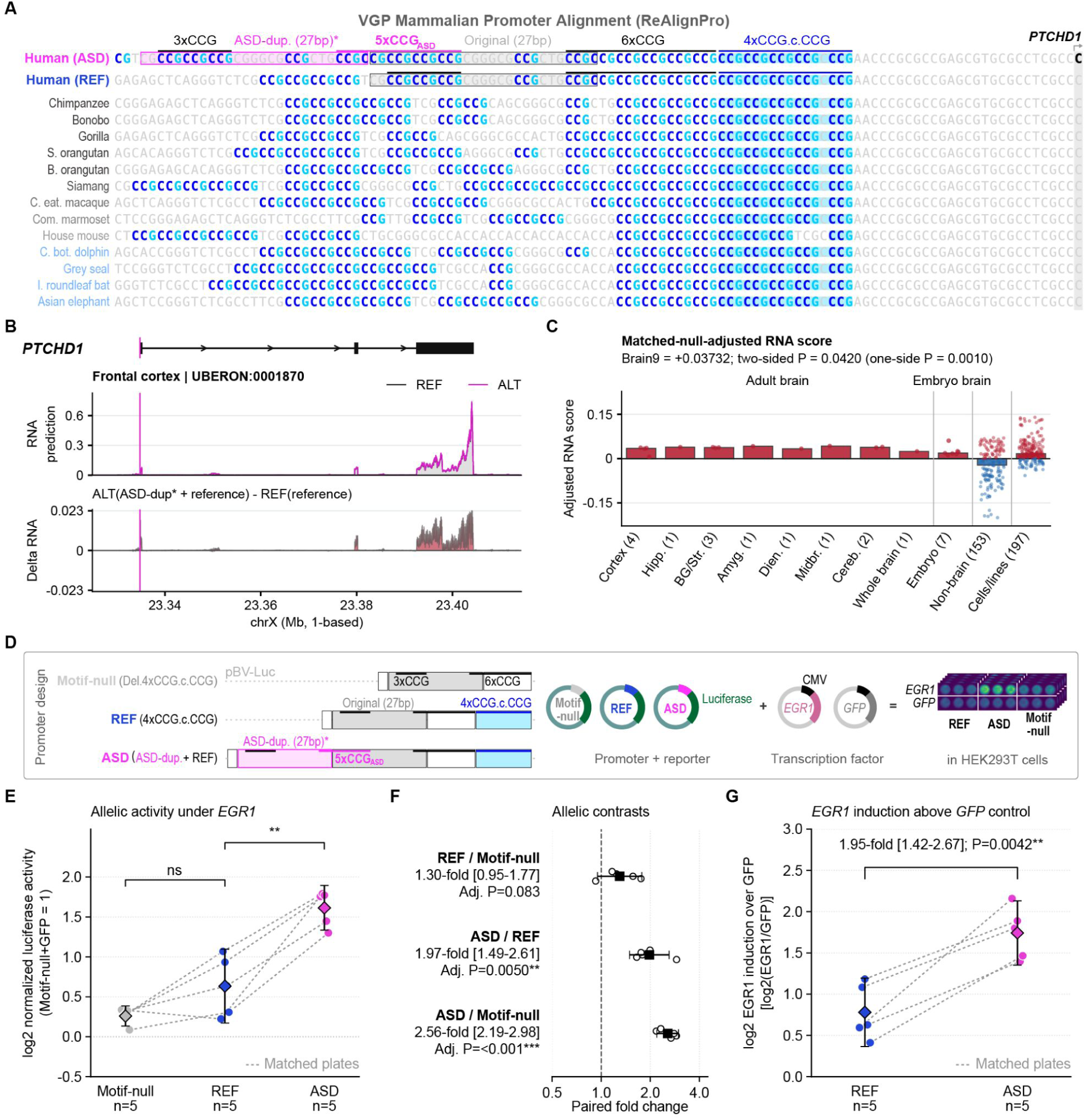
An ASD-associated *PTCHD1* duplication creates an *EGR1*-responsive CCG-repeat gain. (**A**) ReAlignPro alignment of the *PTCHD1* proximal promoter across VGP mammalian species, anchored on the human transcription start site (TSS) and rendered without gaps. A fuller alignment in **Table S2**. Conserved CCG-repeat blocks (3xCCG, 6xCCG) and the *RBFOX1*-like 4xCCG.c.CCG motifs are indicated. The human reference (REF) and the ASD-duplication allele (ASD) are shown on separate rows; the recurrent 27-bp duplication introduces an additional, sequence-distinct 5xCCG run (5xCCG_ASD_) approximately 30 bp upstream of the 4xCCG.c.CCG motif. (**B**) AlphaGenome-predicted whole-brain RNA-seq profiles (UBERON:0000955) for the hg38 reference allele (REF) and the ASD-duplication allele (ALT) in the human genomic context at the *PTCHD1* locus. ΔRNA is defined as ALT minus REF, so that positive values indicate higher predicted output from the ASD allele; note that this is the inverse of the convention in Figures 2B**, C**, and **3D**. (**C**) Matched-null-calibrated AlphaGenome RNA effects summarised by tissue or cell context using Alpha-GENIE (https://alpha-genie.net/#variant). Numbers in parentheses indicate the number of tracks per group. Calibration and grouping procedures are described in **STAR Methods**. (**D**) Luciferase reporters carrying the reference promoter allele (REF, 4xCCG.c.CCG), the ASD-duplication allele (ASD, ASD-dup. + REF), or an engineered motif-null allele (Del.4xCCG.c.CCG) in pBV-Luc, co-transfected into HEK293T cells with CMV-*EGR1* or a CMV-*GFP* expression-vector control. Additional reporter and transcription-factor controls are shown in **Figure S4C,D**. (**E**) Plate-matched promoter activity under *EGR1*. The luciferase signal was background-subtracted, normalized within the plate to motif-null+*GFP*, averaged across technical replicates, and log_2_-transformed. Points represent five independent transfections; dashed lines connect alleles assayed on the same plate, and diamonds with bars show mean log_2_ activity and 95% confidence intervals. Brackets indicate the two sequential adjacent allele comparisons; significance was determined using the same Holm-adjusted P values reported in (**F**). (**F**) Paired allelic fold changes from (**E**). Open circles show plate-level ratios; filled squares and horizontal bars show geometric mean fold changes and 95% confidence intervals. *p-*values were obtained from two-sided paired *t*-tests on log_2_ ratios and adjusted using the Holm method within the pre-specified three-contrast *EGR1* family (m=3). \*\**p* < 0.01. (**G)** *EGR1* induction relative to the *GFP*-vector control, calculated for each allele as log2(*EGR1*/*GFP*). The allele-by-*EGR1* interaction is the difference between ASD and reference induction values, equivalent after back-transformation, and equals (*ASD_EGR1_*/*ASD_GFP_*)/(*REF_EGR1_*/*REF_GFP_*). The interaction *p-*value was obtained from a separate two-sided paired *t-*test and was not included in the Holm family in (**F**). See also **Figure S4** and **Table S2**.

We tested whether promoter duplication affected *PTCHD1* expression and responsiveness to *EGR1*. With the hg38 sequence as REF and the autism-associated 27 bp duplication as ALT (positive ΔRNA = higher predicted output from the duplication allele), AlphaGenome predicted higher *PTCHD1* output from the duplicated allele across all 9 brain tissue groups (**Figure 4B,C**). In contrast, non-brain groups and cells/lines were heterogeneous (**Figure 4B,C**).

To test the prediction that the 27 bp duplication increases promoter activity in ASD patients, we generated reporter constructs containing the hg38 reference promoter of *PTCHD1* (REF), the ASD-duplication promoter of *PTCHD1* (ASD), or the motif-deleted promoter (Del.4xCCG.c.CCG, motif-null). We co-transfected HEK293T cells with these reporters and either CMV-*EGR1* or CMV-*GFP* (**Figure 4D**). Across five independent co-transfections with *EGR1*, luciferase expression from the ASD *PTCHD1* promoter significantly exceeded both REF and motif-null, whereas REF did not significantly exceed the motif-null promoter (**Figure 4E**). The contrast corresponding to the patient variant (ASD vs REF) was 1.97-fold (95% CI, 1.49-2.61; Holm-adjusted P = 0.0050; **Figure 4F**). The contrast between ASD vs motif null was 2.56-fold (95% CI, 2.19-2.98; Holm-adjusted P < 0.001), whereas REF vs motif-null was 1.30-fold and not statistically significant (95% CI, 0.95-1.77; Holm-adjusted P = 0.083; **Figure 4F**).

To determine whether the ASD duplication altered *EGR1* responsiveness, we compared *EGR1*-induced fold activation between the two *PTCHD1* promoter alleles. For each allele, we calculated fold activation by dividing reporter activity under *EGR1* co-transfection by activity under the matched *GFP* control, using that allele’s own control activity as its baseline. This fold activation was 1.95 times higher for the ASD allele than for the REF allele (95% CI, 1.42-2.67; paired P = 0.0042; **Figure 4G**). Thus, the ASD duplication enhanced the *PTCHD1* promoter response to *EGR1*.

The ASD-versus-REF separation was not reproduced under *EGR1ΔBD* (deleted binding domain)*, WT1, WT1ΔBD* (deleted binding domain)*, GFP*, or no added transcription factor, where ASD/REF geometric mean ratios remained near unity (0.99-1.04; **Figure S4C,D**). The patient-variant contrast remained significant after Holm correction, and the separately tested allele-by-*EGR1* interactions. The *PTCHD1* findings therefore provide a patient-derived example in which added CCG copies increase *EGR1*-responsive promoter activity, whereas the human-specific CCG repeat promoter in *RBFOX1* provides an example of a human-specific *EGR1* upregulation that is also impacted in other ASD individuals with a deletion of the CCG repeat promoter.

## Discussion

We asked whether a disease-associated structural variant encompassing an otherwise unique human promoter variant can remove a promoter feature acquired during human evolution. The ASD-risk *RBFOX1* deletion encompasses an alternative promoter and has been associated with reduced *RBFOX1* expression^12,13^. Separately, 17 clinical CNVs reported in 36 individuals with neurodevelopmental phenotypes overlapped the same promoter-containing interval^34^. The human-lineage-unique variant is only a three-base expansion of a CCG promoter repeat. This CCG promoter expansion enhances responsiveness to the *EGR1* transcription factor. We interpret this enhancement as resulting from the CCG expansion creating an additional overlapping human-specific *EGR1* binding site in the *RBFOX1* promoter. Conversely, the ASD-risk *PTCHD1* promoter duplication, which also contains the same CCG repeat sequence, causes increased *EGR1* responsiveness.

We propose that this mechanism connects *RBFOX1* CCG-repeat gains and losses in the human lineage and in ASD individuals, respectively, to the established cis-regulatory roles of short tandem repeats (STRs). Genome-wide studies have linked STR length variation to nearby gene expression^67–70^, while promoter tandem repeats have been implicated in transcriptional evolution^71,72^. STR sequence and copy number can also modulate transcription factor binding^73^. Our reporter assays extend this framework by showing that a single CCG-unit gain increases *EGR1* responsiveness in the *RBFOX1* promoter, connecting human-lineage repeat evolution to a sequence-sensitive regulatory mechanism at a disease-associated locus.

Our results suggest a quantitative potentiation of a pre-existing *EGR1*-responsive promoter element rather than creation of a binary switch. Because *EGR1* is induced by neuronal activity^54,74,75^, the effect of these repeats may itself depend on the activity state of specific brain circuits and their brain regions. Based on our findings and general knowledge of the genes involved, we hypothesize that the human-specific CCG repeat motif in the *RBFOX1* promoter could drive *EGR1* activity-dependent, specialized *RBFOX1* expression in speech brain regions, which would in turn drive differential isoform expression levels of its downstream RNA targets.

We interpret our finding of the same *RBFOX1* CCG repeat promoter sequence in the promoters of 103 other genes as suggesting that the *RBFOX1* promoter belongs to a broad class of promoters. Of these 104 genes, 16 are in the SFARI-annotated gene database, associated with autism^58^, and are enriched for nervous system development and postsynaptic compartments. But only a subset of these promoters are predicted (according to AlphaGenome^61^) to regulate their genes in the same way as *RBFOX1*, suggesting that the repeat is not a sufficient portable activating module, but that local sequence and genomic context likely modulate its effect^76,77^. *AUTS2* emerged as a candidate because its predicted brain-effect profile resembled that of *RBFOX1*; *AUTS2* has been disrupted by a *de novo* translocation in an individual with autism^78^. Both *RBFOX1* and *AUTS2* genes were downregulated in the same layer 6 excitatory-neuron cluster in the ASD cortex^59^.

*PTCHD1*^62–66^ emerged as another candidate, with an ASD patient-associated promoter repeat gain. Its reference CCG-repeat architecture is ancestral, whereas a recurrent 27-bp promoter duplication reported in three individuals with ASD^33^ adds an uninterrupted 5xCCG module. The allele-by-*EGR1* interaction indicates that the promoter duplication in ASD individuals preferentially increased *EGR1* responsiveness rather than simply recapitulating an allelic difference present under the *GFP*-vector control. *PTCHD1* therefore provides an independent example of a gain-of-repeat that complements the human-lineage *RBFOX1* insertion and strengthens the broader conclusion that short CCG-repeat gains can tune *EGR1*-responsive promoter output in distinct local contexts. The only previous functional test of this duplication reported a 26% reduction in basal activity from a 3,078-bp upstream fragment, with no significant difference with its 200-bp construct^33^; our core-promoter reporters likewise showed no allelic difference with the *GFP*-vector control.

Our findings have been made possible by high-quality genome assemblies from the Vertebrate Genomes Project^27,40^ and its collaborative consortia^29–31^, which now resolve repeat-rich promoters at base-pair resolution. Our ReAlignPro software enabled promoter-anchored comparative alignments to identify and visualize repeat-copy changes that substitution-rate screens miss. Calibrated variant-effect models and reporter assays across species and patient sequences provide evidence-driven functional hypotheses. Each biological level of analysis answered a different question: what changed, which effect direction is predicted, and whether the variant alters promoter activity. This framework can be used in future studies of genes with regulatory specializations and transcription factor responsiveness.

More broadly, this study connects a clinically observed genomic promoter micro-deletion to a human-specific promoter and a *de novo* molecular function. As many promoters contain CGG repeats, such variation has the opportunity to arise in many places, perhaps contributing to the diverse pathophysiology within behaviorally defined disorders such as ASD. Although genomic insertion/deletion in repeat-rich promoters has resulted in assembly problems in the past, they are now tractable experimental substrates for studying human regulatory evolution and disease-associated variations. Together, these results demonstrate the utility of modern genomic technologies and analyses in resolving promoter sequence variation and highlight the role of promoter CCG repeat sequence content in tuning transcriptional responses to *EGR1*.

### Possible limitations of the study

Our conclusions are limited in several ways. AlphaGenome annotation does not infer gene expression in activity-dependent neuronal states or across individual genetic backgrounds, while promoter reporter constructs in HEK293T cells do not account for neuronal chromatin, long-range regulation, endogenous factor concentrations, direct occupancy, or specialized gene regulation in speech/song brain areas. These limitations can be addressed through isogenic editing in the human brain, cultured neurons, or cortical organoids; activity-dependent perturbation of *EGR1;* promoter-resolved CAGE or long-read isoform measurements; and direct occupancy assays. The clinical variants in *RBFOX1* and *PTCHD1* do not isolate the CCG repeat effects from the surrounding sequence, so matched constructs that preserve variant length and GC content while selectively verifying the deleted/added CCG repeat will be required to isolate CCG-repeat-specific activity. Even when all such experiments are conducted, we don’t see them changing the major findings of this study: that a small CCG promoter repeat addition in the human lineage can change the regulation of the gene and that this same region has structural variants in some autistic individuals that are also associated with a change in gene regulation.

## Supporting information

STAR Methods

## Acknowledgments

We thank all members of the VGP, T2T, and HPRC consortia. We thank the NCBI eukaryote annotation team, the Ensembl annotation team, the ENCODE consortium, the Zoonomia consortium, the JASPAR database, the AADR database, the NCBI Sequence Read Archive, dbSNP, and PsychENCODE for public data, and the UCSC Genome Browser and WashU Comparative Epigenome Browser for public data browsers. This study uses data generated by the DECIPHER community; a full list of contributing groups is available at https://deciphergenomics.org and by email at. We thank Drs. P. Rajasethupathy, C. Scharff, and M. Marwa for scientific support and comments on an earlier version of this work; N. Desilva and Dr. M. Biegler for help establishing HEK293T cell culture; and Z. Zargoozian and S. Bollu for help cloning the ΔBD plasmids. This work was supported by NIH grant R01DC018691 (to C.L. and E.D.J.), an NSF Graduate Research Fellowship (to M.H.D.), and the Howard Hughes Medical Institute (to M.H.D. and E.D.J.).

## Author contributions

CL designed the multi-omic analyses, collected datasets, developed ReAlignPro and Alpha-GENIE, conducted all computational analyses, and built the UCSC TrackHub. MHD identified *RBFOX1* as a gene of interest. CL identified human-specific CCG repeats in the promoters of ASD-risk genes *RBFOX1* and *PTCHD1* as regulatory elements of interest. CL and MHD designed the luciferase reporters for *RBFOX1.* CL designed the ASD-related luciferase reporters for *PTCHD1*. MHD and CL designed the TF vectors and Δ_BD_ plasmids. CL conducted luciferase assays. CL wrote the draft paper. All authors reviewed the draft. EDJ secured funding and supervised the study.

## Declaration of generative AI and AI-assisted technologies in the writing process

During the preparation of this work, the authors wrote the first draft of the manuscript without any generative AI or AI-assisted technologies. Afterward, the authors used *OpenAI ChatGPT* (versions 5.6 Solo and 6.0 Astra) to improve the manuscript’s readability and language, and used *Grammarly* to check grammar and typos. The authors used *OpenAI Codex* (versions 5.5, 5.6, and 6.0) and *Anthropic Claude code* (versions Opus 5 and Fable 5.1) to develop and debug the source code for ReAlignPro and Alpha-GENIE. After using these tools, the authors reviewed and edited the content as needed and took full responsibility for the content of the publication.

## Inclusion and diversity

We support inclusive, diverse, and equitable research practices.

