## Supplementary Information for "Human-specific CCG-repeat insertion in the *RBFOX1* promoter implicated in autism and speech enhances *EGR1*-responsive activity"

**This PDF file includes:**

Figures S1-4

Table legends S1-4 (GoogleSheetLink)

Supplementary References

### Supplemental information

#### Supplementary Figures

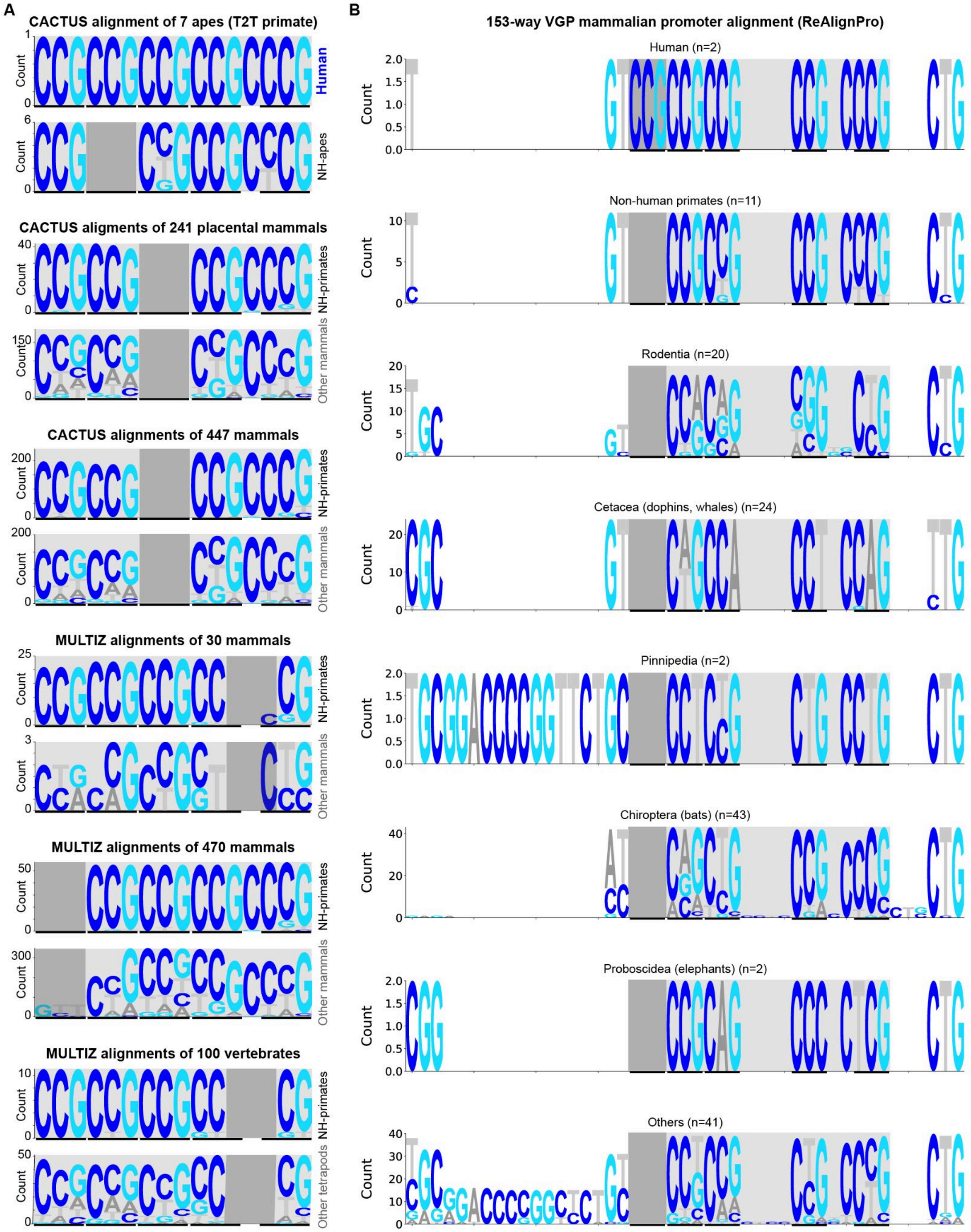

**Figure S1. CCG-repeat architecture of the homologous *RBFOX1* core promoter regions across mammals, related to Figure 1.**

(**A**) Sequence logos summarizing the same *RBFOX1* repeat interval from independent whole-genome alignment resources, including the 8-way T2T primate CACTUS alignment[31], 241-way placental-mammal or 447-way broader mammalian CACTUS alignments, and MULTIZ alignments of 30 mammals, 470 mammals, and 100 vertebrates in the UCSC genome browser[41]. For each alignment resource, logos are shown separately for human or non-human primates/apes, as indicated, and for a comparison group of other mammals or other tetrapods. The shaded pink region denotes the homologous repeat block, and dark pink blocks indicate homologous alignment columns of the human-specific trinucleotide insertion that are absent, gapped, or poorly resolved in a given resource. Across assemblies and alignment strategies, the repeat-rich context yields alternative phasing and gap placement around the tract. Still, it consistently supports an expanded CCG-repeat configuration in humans relative to non-human primate orthologs.

(**B**) Sequence logos summarizing the aligned *RBFOX1* core-promoter interval containing the human-specific trinucleotide insertion across 144 mammalian assemblies from the Vertebrate Genomes Project (n = 145, including hg38). Assemblies are grouped by major lineages, including humans, non-human primates, Rodentia, Cetacea, Pinnipedia, Chiroptera, Proboscidea, and others, with sample sizes indicated for each group. The shaded pink region marks the homologous regions of the human CCG-repeated motif (4xCCG.c.CCG) within the shared alignment. Letter height reflects nucleotide counts at each aligned position (y-axis).

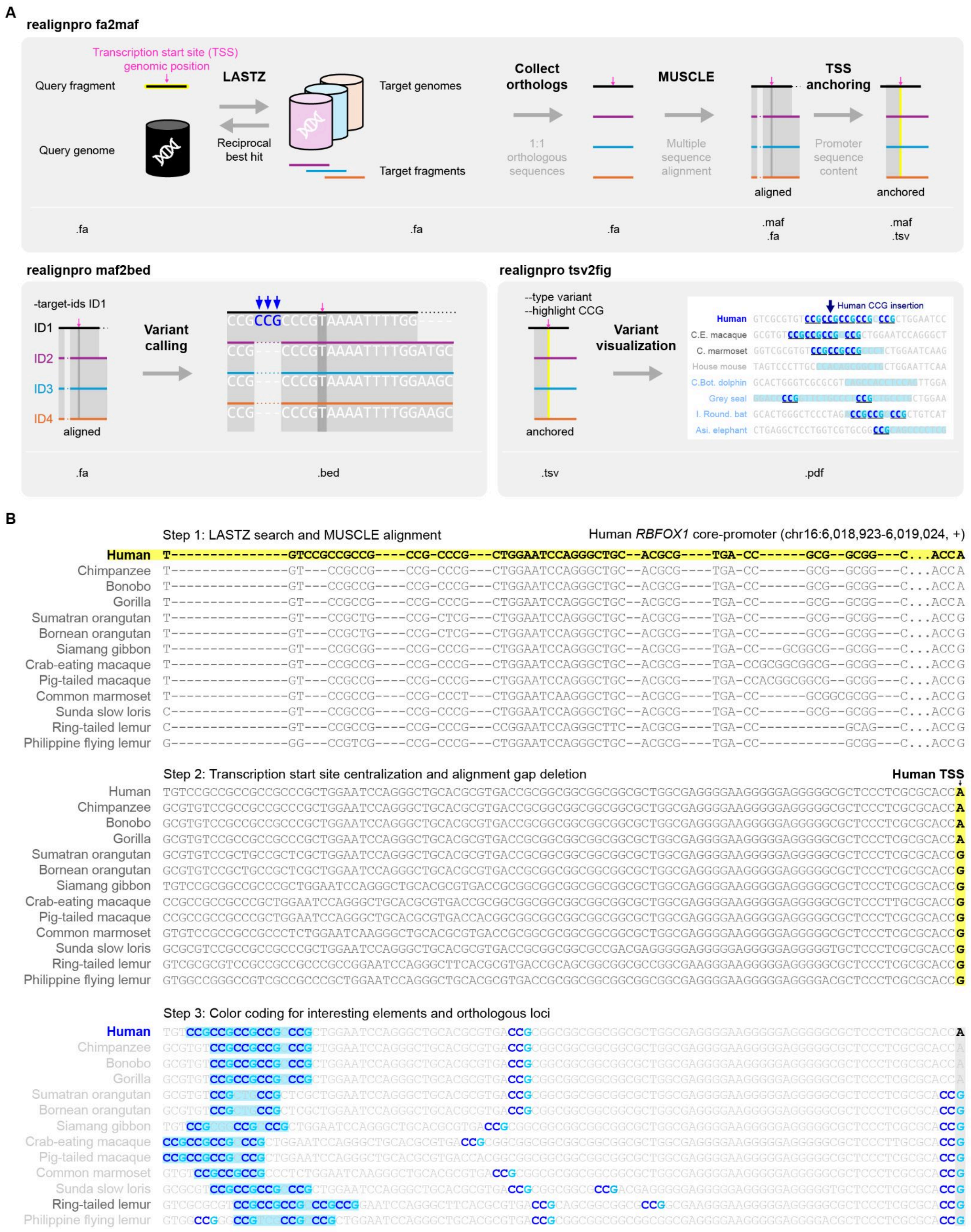


**Figure S2. ReAlignPro workflow and visualization output, related to Figure 1.**

(**A**) Re-Align Promoter (ReAlignPro) workflow and associated functions. Given a query promoter fragment, the query genome, and the genomic position of the reference transcription start site (TSS), *realignpro fa2maf* identifies one-to-one orthologous sequences in target genomes using LASTZ searches and reciprocal-best-hit filtering. These sequences are aligned using MUSCLE, followed by alignment-gap removal and anchoring to the reference TSS. Aligned sequences are exported in FASTA and MAF formats, and TSS-anchored sequences in MAF and TSV formats. *realignpro maf2bed* performs variant calling from the aligned sequences and exports results in BED format. *realignpro tsv2fig* generates PDF visualizations from the TSS-anchored TSV output, with options for variant visualization and sequence-motif highlighting, illustrated here using CCG.

(**B**) ReAlignPro alignment and TSS-anchored visualization of the RBFOX1 core promoter interval across 13 species of Primatomorpha from the 145-genome VGP mammalian dataset. Step 1 shows the multiple-sequence alignment generated following LASTZ searches and MUSCLE alignment, with the human reference promoter sequence highlighted in yellow. Step 2 removes alignment gaps and anchors each sequence to the position corresponding to the human TSS, highlighted in yellow, producing a gapless sequence map that facilitates interpretation in this repeat-rich region. In step 3, CCG motifs are highlighted in bold blue and sky blue, and pale-blue shading marks the repeat-containing region and its corresponding regions across species. Unhighlighted nucleotides are shown in light grey, and the column corresponding to the human TSS is shaded grey. Full sequence outputs for all three steps are provided in **Table S2**.

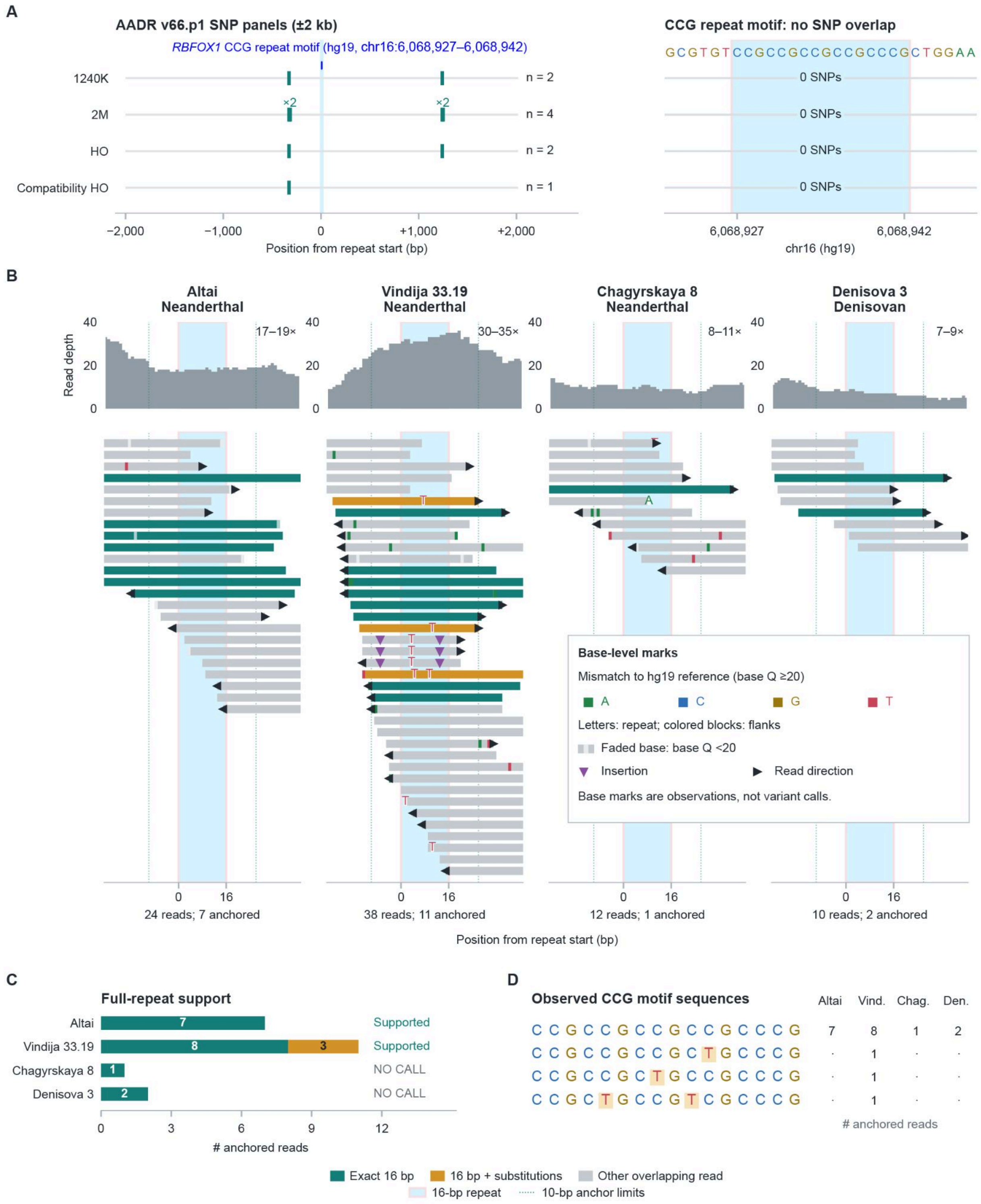


65

66 **Figure S3. AADR SNP panel coverage and archaic read support at the *RBFOX1* CCG
67 repeat motif, related to Figure 1.**

(**A**) Positions of SNPs included in the AADR v66.p1 1240K, 2M, Human Origins (HO), and compatibility HO panels around the *RBFOX1* CCG repeat motif (CCGCCGCCGCCGcCCG, 4xCCG.c.CCG; hg19/b37 chr16:6,068,927–6,068,942). The left view shows the repeat and 2-kb flanks; teal ticks indicate panel SNP positions, and n denotes the number of sites within the displayed interval. The right view shows the reference sequence and an enlarged view of the CCG repeat motif interval. None of the four panels contains a SNP within the CCG repeat motif. These tracks represent panel marker locations; absence of a marker does not establish absence of variation or confirm the reference repeat allele.

(**B**) Read depth and BAM alignments for three Neanderthals—Altai, Vindija 33.19, and Chagyrskaya 8—followed by Denisova 3. Each view spans the repeat and 25-bp flanks. Primary, non-duplicate, QC-passing alignments with mapping quality ≥ 30 were retained; MAPQ 255 was excluded. Coverage includes A/C/G/T observations with base quality ≥ 20 and is displayed on a common 0–40 scale. Each read-track row represents an alignment that overlaps the CCG repeat motif, without downsampling. Coverage outside the repeat can additionally include reads that do not overlap the motif. Skyblue shading marks the repeat, and dotted lines delimit the outer boundaries of its 10-bp anchors. Teal reads contain the exact sequence ‘CCGCCGCCGCCGcCCG’ and both exact anchors; orange reads satisfy the same anchoring criteria but contain substitutions within a 16-bp sequence. Both categories require base quality ≥ 20 throughout the anchors and intervening sequence. Gray reads do not satisfy the complete anchored-read criteria. Reference-mismatching bases are shown as letters within the repeat and colored blocks in the flanks: A, green; C, blue; G, gold; and T, red. Lower-quality bases are faded. Purple triangles indicate insertions, and black arrowheads indicate alignment direction at visible read ends. These base-level marks represent observations, not independently established variant calls.

(**C**) Full-repeat support after grouping reads by library, read identifier, and fragment endpoints into putative molecule groups. Exact 16-bp support comprises 7, 8, 1, and 2 groups for Altai, Vindija, Chagyrskaya, and Denisova, respectively. Vindija additionally contains three 16-bp observations with substitutions. "Supported" requires at least three exact-support groups without fully anchored alternative-length or conflicting observations. Chagyrskaya and Denisova remain "NO CALL" because support falls below this threshold; this does not indicate absence of the repeat.

(**D**) Distinct 16-bp sequences observed among passing molecule groups and their counts in each individual. Substitutions relative to the reference repeat are highlighted, and dots indicate zero observations. Counts summarize sequence evidence under the specified filters and do not establish diploid genotypes or population allele frequencies.

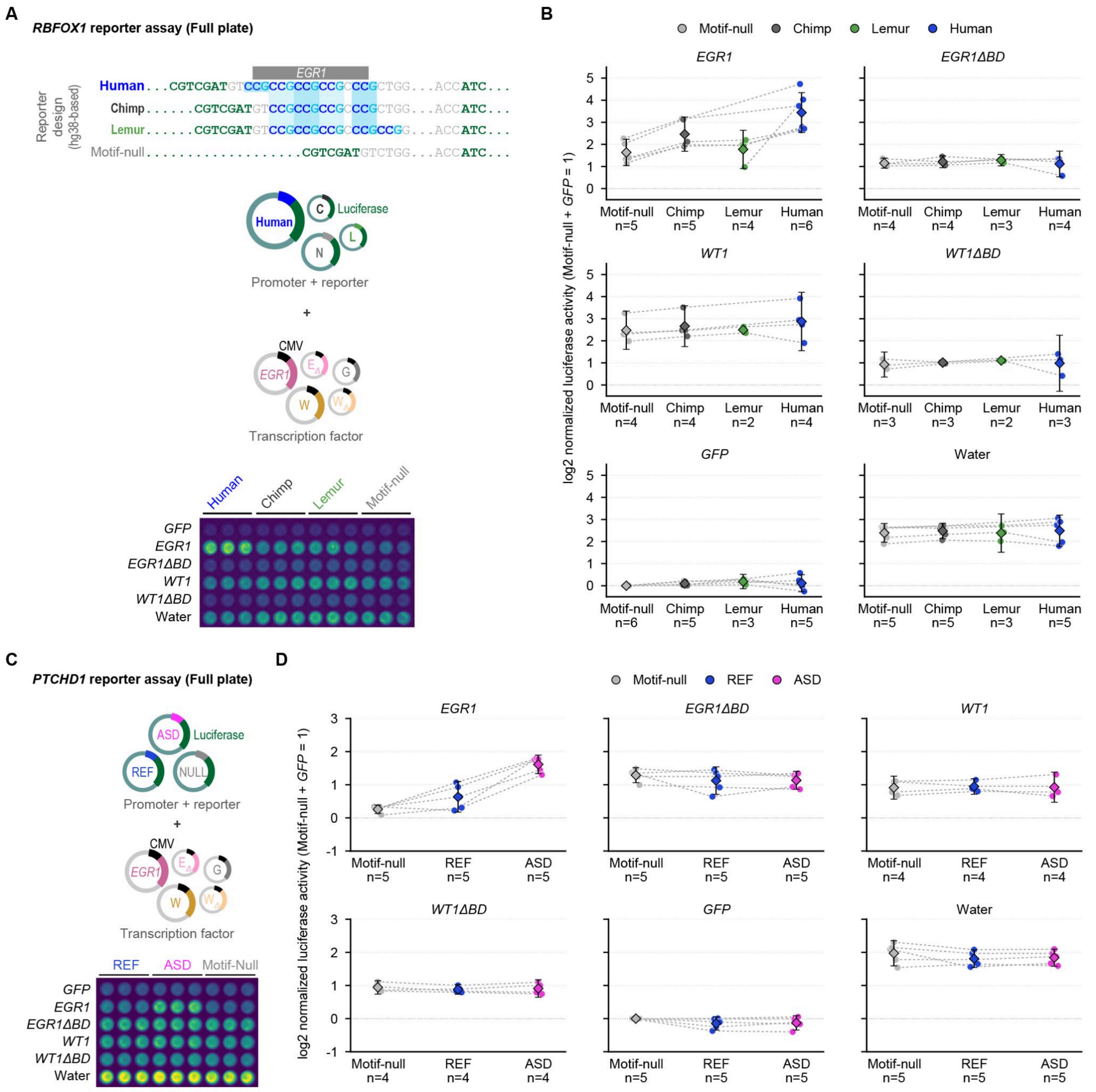


**Figure S4. Reporter designs, representative assay plates, and complete *RBFOX1* and *PTCHD1* reporter matrices, related to Figures 2 and 4.**

(**A**) *RBFOX1* reporter assay design. Matched promoter fragments carrying the motif-null, chimpanzee, ring-tailed lemur, or human allele were cloned upstream of luciferase and co-transfected into HEK293T cells with CMV-driven *EGR1, EGR1ΔBD, WT1, WT1ΔBD,* or

*GFP* expression plasmids, or with a water-only control lacking an added expression plasmid. ΔBD denotes deletion of the zinc-finger DNA-binding domain. In the plasmid schematics, C, L, and N denote the chimpanzee, lemur, and motif-null reporters, respectively; EΔ, W, WΔ, and G denote *EGR1ΔBD, WT1, WT1ΔBD,* and *GFP*, respectively. The representative luminescence image shows the plate layout for the reporter-by-expression condition.

(**B**) Complete *RBFOX1* reporter matrix across *EGR1, EGR1ΔBD, WT1, WT1ΔBD, GFP*, and water-control conditions.

(**C**) *PTCHD1* reporter-assay design. Matched promoter fragments carrying the motif-null, reference (REF), or ASD-duplication (ASD) allele were cloned upstream of luciferase and tested using the same expression-plasmid and control conditions as in (A). The representative luminescence image shows the corresponding reporter-by-expression-condition plate layout.

(**D**) Complete *PTCHD1* reporter matrix across *EGR1, EGR1ΔBD, WT1, WT1ΔBD, GFP*, and water-control conditions.

For (**B**) and (**D**), luminescence signals were background-subtracted using no-reporter wells and normalized within each plate to the background-subtracted motif-null+GFP condition before $\log_2$ transformation. Three technical wells per condition were averaged within each independent transfection experiment. Each point represents one independent transfection; dashed lines connect reporter alleles measured on the same plate under the same expression condition. Diamonds indicate mean $\log_2$ activity, and error bars indicate 95% confidence intervals of the mean when $n \geq 3$. The n shown beneath each reporter–condition combination denotes the number of independent transfections. Conditions with fewer than three independent transfections are presented descriptively without inferential interpretation. The plate images in (**A**) and (**C**)

131 illustrate assay layout and relative luminescence within a representative plate. Primary allelic
132 contrasts and allele-by-*EGR1* interaction tests are reported in **Figures 2E–G** and **4E–G**.

#### Supplementary Tables

**Table S1. The autism-risk deletion (AU077504) and the DECIPHER clinical copy number variants overlapped with the autism-risk *RBFOX1* promoter, as shown in Figure 1.**

**Table S2. 153-way VGP mammalian genomes for ReAlignPro alignments of *RBFOX1* and *PTCHD1*, related to Figures 1 and 4.**

**Table S3. Transcription factor binding motif estimations of homologous loci of the human 4xCCG.c.CCG motif, related to Figure 2.**

**Table S4. *RBFOX1*-like genes with the 4xCCG.c.CCG motif in core promoters, as shown in Figure 3.**
