## Supplementary material for "Human-specific CCG-repeat insertion in the *RBFOX1* promoter implicated in autism and speech enhances *EGR1*-responsive activity": STAR Methods

### STAR Methods for

5 Lead contact

Supplementary Tables: GoogleSheetLink

### RESOURCE AVAILABILITY

#### Lead contact

#### Materials availability

This study generated *RBFOX1* and *PTCHD1* promoter–luciferase reporters and *WT1, GFP,* and *EGR1/WT1* DNA-binding-domain-deletion expression constructs (https://github.com/chulbioinfo/alphagenie).

#### Data and code availability

The public genome assemblies, annotations, and sequencing resources used in this study are identified in the key resources table and in the relevant STAR Methods sections. ReAlignPro source code is available at https://github.com/chulbioinfo/ReAlignPro, and the UCSC track hub is available at https://ccg2025.s3.us-east-2.amazonaws.com/hub.txt.

AlphaGENIE source code, installation instructions, analysis documentation, and verification reports are publicly available under release v0.24.0. The release includes variant definitions, matched-null designs, track assignments, processed source tables, group and null-consensus summaries, brain matrices, saved figures, and frontal cortex curve artifacts. Full canonical track-level real/null arrays are included for the standalone *RBFOX1* and *PTCHD1* analyses; the 17-variant package contains Brain9 null arrays but not the complete raw track-level null outputs for every candidate. Private raw jobs from the fresh verification run and reference genomes are not bundled. The saved manuscript results can be inspected and downloaded through the public explorer (https://alpha-genie.net).

### STAR METHODS

#### KEY RESOURCES TABLE

| REAGENT or RESOURCE | SOURCE | IDENTIFIER |
|---|---|---|
| **Chemicals, peptides, and recombinant proteins** | | |
| Lipofectamine 2000 | Thermo Fisher Scientific | Cat#11668019 |
| Opti-MEM | Gibco | Cat#31985070 |
| Fetal bovine serum | Gibco | Cat#A5670701 |
| Acridine orange/propidium iodide stain | Logos Biosystems | Cat#F23001 |
| **Critical commercial assays** | | |
| ONE-Glo Luciferase Assay System | Promega | Cat#E6110 |
| HiFi DNA Assembly reagents | New England Biolabs | Cat#E5520S |
| GeneArt Complex Gene Synthesis | Thermo Fisher Scientific | ProjectID: 2025AAN2TD / 2025ABMY6D |
| **Deposited data** | | |
| Human GRCh38.p14 / hg38 genome | NCBI RefSeq | GCF_000001405.40 (https://www.ncbi.nlm.nih.gov/datasets/genome/GCF_000001405.40/) |
| Rhesus macaque Mmul_10 / rheMac10 genome | NCBI RefSeq | GCF_003339765.1 (https://www.ncbi.nlm.nih.gov/datasets/genome/GCF_003339765.1/) |
| Common marmoset calJac4 genome | NCBI RefSeq | GCF_009663435.1 (https://www.ncbi.nlm.nih.gov/datasets/genome/GCF_009663435.1/) |
| VGP Phase I v1.0 mammalian assembly set | Formenti et al.[1] | 153 input assemblies; accession manifest in Table S2: https://github.com/VGP/vgp-phase1 |
| 14-way Primatomorpha CACTUS alignment | This study | https://ccg2025.s3.us-east-2.amazonaws.com/data/maf/14primates_hg38.bb |
| Independent CACTUS/MULTIZ alignment resources | UCSC Genome Browser[2]; Yoo et al.[3] | https://cglgenomics.ucsc.edu/february-2024-t2t-apes/ http://hgdownload.soe.ucsc.edu/goldenPath/hg38/cactus241way/ http://hgdownload.soe.ucsc.edu/goldenPath/hg38/cactus447way/ |

| | | |
|---|---|---|
| | | http://hgdownload.soe.ucsc.edu/goldenPath/hg38/multiz30way/ http://hgdownload.soe.ucsc.edu/goldenPath/hg38/multiz470way/ http://hgdownload.soe.ucsc.edu/goldenPath/hg38/multiz100way/ |
| Clinical *RBFOX1* variants | Sebat et al.[4]; Martin et al.[5]; DECIPHER[6] | AU077504; https://www.deciphergenomics.org/; This study Table S1 |
| HAR coordinate catalog | Cui et al.[7] | Cui et al. Supplementary Table 2 |
| HAQER coordinate catalog | Mangan et al.[8] | https://vertgenlab.cloud.duke.edu/publications/haqer/regionsOfInterest/haqersByAssembly/haqer.hg38.bed |
| FANTOM5 CAGE track | Forrest et al.[9] | FANTOM5 reprocessed7 https://hgdownload.soe.ucsc.edu/gbdb/hg38/fantom5/hg38.cage_peak.bb |
| ENCODE cCRE track | Moore et al.[10] | Version 4; https://hgdownload.soe.ucsc.edu/gbdb/hg38/encode4/ccre/encodeCcreRegistry.bb |
| Primary motor cortex single-nucleus RNA-seq tracks | Comparative Epigenome Browser[11] | https://epigenome.wustl.edu/renlab/human_m1_rna/ https://epigenome.wustl.edu/renlab/macaque_m1_rna/ https://epigenome.wustl.edu/renlab/marmoset_m1_rna/ |
| rheMac10-hg38 chain | UCSC Genome Browser[2] | https://hgdownload.soe.ucsc.edu/goldenPath/rheMac10/liftOver/rheMac10ToHg38.over.chain.gz |
| calJac4-hg38 chain | UCSC Genome Browser[2] | https://hgdownload.soe.ucsc.edu/goldenPath/calJac4/liftOver/calJac4ToHg38.over.chain.gz |
| Arcseqhub | Liang et al.[12] | https://hgdownload.soe.ucsc.edu/gbdb/hg38/ancient/neanderthal.hg38.filt.vcf.gz https://hgdownload.soe.ucsc.edu/gbdb/hg38/ancient/denisovan.hg38.filt.vcf.gz |
| The Allen Ancient DNA Resource (AADR) | Mallick et al.[13] | v66.1 https://dataverse.harvard.edu/dataset.xhtml?persistentId=doi:10.7910/DVN/FFIDCW |
| Altai (Denisova 5), chromosome 16 | Prüfer et al.[14]; MPI-EVA | https://cdna.eva.mpg.de/neandertal/altai/AltaiNeandertal/bam/AltaiNea.hg19_1000g.16.dq.bam |
| Vindija 33.19, chromosome 16 | Prüfer et al.[15]; MPI-EVA | https://cdna.eva.mpg.de/neandertal/Vindija/bam/Pruefer_etal_2017/Vindija33.19/Vi33.19.chr16.indel_realn.bam |

| | | |
|---|---|---|
| Chagyrskaya 8, chromosome 16 | Mafessoni et al.[16]; MPI-EVA | https://cdna.eva.mpg.de/neandertal/Chagyrskaya/BAM/chr16-reali.bam |
| Denisova 3, whole-genome BAM | Meyer et al.[17]; MPI-EVA | https://cdna.eva.mpg.de/denisova/alignments/T_hg19_1000g.bam |
| Denisova 3, alternative processed chr16 BAM | MPI-EVA; same individual as above | Processing-sensitivity analysis only https://cdna.eva.mpg.de/neandertal/Vindija/bam/Pruefer_etal_2017/Denisova3_realigned/chr16-reali.bam |
| HPRC haplotype pangenome alignment | Liao et al.[18] | https://s3-us-west-2.amazonaws.com/human-pangenomics/index.html?prefix=pangenomes/freeze/freeze1/minigraph-cactus/hprc-v1.1-mc-grch38/ |
| dbSNP155 | Sherry et al.[19] | https://hgdownload.soe.ucsc.edu/gbdb/hg38/snp/dbSnp155Common.bb |
| ClinVar | Landrum et al.[20] | https://hgdownload.soe.ucsc.edu/gbdb/hg38/snp/dbSnp155ClinVar.bb |
| JASPAR CORE 2022 vertebrate matrices | Castro-Mondragon et al.[21] | JASPAR2022_CORE_vertebrates_non-redundant_pfms_meme (https://jaspar.elixir.no/download/data/2026/CORE/JASPAR2026_CORE_vertebrates_non-redundant_pfms_meme.zip) |
| Human gene annotation for AlphaGENIE local configuration | GENCODE[22] | Release 46 GTF (https://www.gencodegenes.org/human/release_46.html); MANE Select flags used when present |
| SFARI Gene 2025 Q4 | SFARI Gene[23] | https://s3.amazonaws.com/sfari-gene/wp-content/uploads/2026/01/14213706/Q4_2025_Report.pdf |
| PsychENCODE ASD frontal-cortex expression resource | Wamsley et al.[24] | http://solo.bmap.ucla.edu/asdscgene/ |
| Reporter measurements | This study | https://github.com/chulbioinfo/alphagenie/assays/ |
| AlphaGENIE manuscript prediction artifacts | This study | Scientific source v0.20 (https://github.com/chulbioinfo/alphagenie/tree/v0.24.0/data/manuscript_v020_20260909) |
| Brain9 track classification | This study | Brain9_adult8_embryo_v1_20260909 (https://github.com/chulbioinfo/alphagenie/blob/v0.24.0/docs/BRAIN9.md) |
| **Experimental models: Cell lines** | | |
| HEK293T | Cellosaurus | RRID: CVCL_0063 (Female) |
| **Bacterial and virus strains** | | |

| | | |
|---|---|---|
| NEBuilder® HiFi DNA Assembly Cloning Kit, With Competent Cells | New England Biolabs | Cat#E5520S |
| **Oligonucleotides** | | |
| Promoter oligonucleotides, *WT1*/*GFP* gBlocks, ΔBD primers | IDT; this study | https://github.com/chulbioinfo/alphagenie/assays/ |
| **Recombinant DNA** | | |
| pBV-Luc | Addgene | Plasmid#16539 |
| pcDNA3-*EGR1* | Addgene | Plasmid#11729 |
| *RBFOX1* and *PTCHD1* promoter–luciferase reporters | This study | https://github.com/chulbioinfo/alphagenie/assays/ |
| pcDNA3-*WT1*, pcDNA3-*GFP*, pcDNA3-*EGR1ΔBD*, pcDNA3-*WT1ΔBD* | This study | https://github.com/chulbioinfo/alphagenie/assays/ |
| **Software and algorithms** | | |
| ReAlignPro | This study | https://github.com/chulbioinfo/ReAlignPro |
| AlphaGenome and Python API client | Avsec et al.[25] | SDK 0.8.0 (https://github.com/google-deepmind/alphagenome) |
| AlphaGENIE analysis and visualization | This study | https://alpha-genie.net, https://github.com/chulbioinfo/alphagenie/ |
| Python environment for AlphaGENIE v0.24 verification | Python Software Foundation and package developers | Python 3.11; NumPy 2.4.6; pandas 2.3.3; Matplotlib 3.11.1; locked dependencies (https://github.com/chulbioinfo/alphagenie/blob/v0.24.0/requirements-lock.txt) |
| LASTZ | Harris[26] | v1.04.52 |
| MUSCLE | Edgar[27] | v3.8.1551 |
| samtools | Li et al.[28] | v1.21 |
| NCBI Datasets | O'Leary et al.[29] | v16.41.0 |
| CrossMap | Zhao[30] | v0.7.0 |
| BEDTools | Quinlan and Hall[31] | v2.31.1 |
| BLAST | Ye et al.[32] | v2.16.0+ |
| ROADIES | Gupta et al.[33] | Main branch commit 6447610 at 2025-05-27 between v0.1.10 and v0.2.0 https://github.com/TurakhiaLab/ROADIES |

| | | |
|---|---|---|
| Progressive Cactus | Armstrong et al.[34] | CACTUS v3.2.0; toil v9.3.0; https://github.com/ComparativeGenomicsToolkit/cactus |
| Logomaker | Tareen et al.[35] | v0.8.7; https://logomaker.readthedocs.io/en/latest/ |
| FIMO | Grant et al.[36] | v5.5.9 |
| g:Profiler | Reimand et al.[37] | https://biit.cs.ut.ee/gprofiler/gost (1-18-2026) |
| FIJI/ImageJ | Schindelin et al.[38] / Abràmoff et al.[39] | v2.16.0 |
| UCSC Genome Browser | Perez et al.[2] | https://genome.ucsc.edu |
| WashU Comparative Epigenome Browser | Zhuo et al.[40] | https://epigenome.wustl.edu/BrainComparativeEpigenome/ |

#### STUDY PARTICIPANT AND EXPERIMENTAL MODEL DETAILS

##### Public genomic and transcriptomic resources

Comparative analyses used publicly available modern human and non-human animal genome assemblies and associated annotations, including resources from the Vertebrate Genomes Project (VGP)[1,41] and collaborating consortia[3,18,42]. Clinical analyses used previously reported *RBFOX1*[4,5] and *PTCHD1*[43] variants and DECIPHER annotations[44]. Archaic hominin resources included ArcSeqHub variant catalogs[12], SNP-marker panels from the Allen Ancient DNA Resource (AADR)[13], and published high-coverage genomic read alignments from the Max Planck Institute for Evolutionary Anthropology (MPI-EVA)[14–17]. Independent cross-species whole-genome alignments were obtained from the telomere-to-telomere (T2T) ape project[3] and the UCSC Genome Browser[2].

Cell-type-specific single-nucleus RNA-seq coverage tracks[11] for the primary motor cortex of humans, rhesus macaques, and common marmosets were obtained from the WashU Comparative Epigenome Browser[40]. Cell-subtype-specific ASD-versus-control differential expression statistics from the PsychENCODE frontal cortex single-nucleus RNA-seq resource were obtained from the ASD Single-Cell Gene Expression Portal (ASD scGENE Portal)[24].

The provenance and processing of these datasets are described below. This study generated no new human genomic or transcriptomic data.

#### HEK293T cells

Reporter assays were performed in HEK293T cells (RRID: CVCL_0063; passages 5-15) maintained at 37°C with 5% $CO_2$ in DMEM (Corning, MT10027CV) supplemented with 10% fetal bovine serum (Gibco, A5670701) and 1% penicillin–streptomycin (Gibco, 15140122). Cells were used at approximately 90–95% confluence before transfection.

### METHOD DETAILS

#### Clinical variants and promoter annotations at *RBFOX1*

The previously reported AU077504 deletion affecting an alternative *RBFOX1* promoter was mapped to hg38 chr16:6,002,834-6,210,814[4,5]. To obtain these coordinates, 201-bp sequences at each breakpoint of the hg18 interval chr16:5,992,836-6,200,816 were extracted and mapped to hg38 using UCSC BLAT[45]. Clinical copy-number variants overlapping the promoter-containing interval were curated from DECIPHER[44], along with the reported phenotype annotations. The current Results summarize 17 CNVs in 36 individuals (**Figure 1A**; **Table S1**). The publicly accessible session is https://genome.ucsc.edu/s/clee03/CCG2026_Fig1A_ASDdel in the UCSC Genome Browser[2].

FANTOM5 CAGE peaks[9] and ENCODE candidate cis-regulatory elements (v4)[10] were used to annotate transcription-initiation and regulatory features at the alternative *RBFOX1* promoter (**Figure 1B**).

**Overlap with human sequence-acceleration catalogs**

The *RBFOX1* promoter and motif-bearing core-promoter intervals were assessed for overlap with published human accelerated regions (HARs)[7,46] and human ancestor quickly evolved regions (HAQERs)[7,8]. The HAR catalog contained 3,257 nonredundant loci from **Supplementary Table 2** of Cui et al.[7]. The HAQER catalog contained 1,581 loci from the hg38 BED file provided by Mangan et al.[8] ( https://vertgenlab.cloud.duke.edu/publications/haqer/regionsOfInterest/haqersByAssembly/haqer.hg38.bed). Sorted hg38 BED intervals were intersected separately with each catalog using BEDTools v2.31.1. Overlap was defined as at least 1 shared base pair, with no extension of the input intervals. The positions of HARs and HAQERs relative to the *RBFOX1* gene body are shown in **Figure 1A**.

**Cross-species M1C snRNA coverages of human, macaque, and marmoset**

Human (GRCh38.p14/hg38; GCF_000001405.40), rhesus macaque (Mmul_10/rheMac10; GCF_003339765.1), and common marmoset (calJac4; GCF_009663435.1) reference assemblies were used for the published primary motor cortex single-nucleus transcriptome study[11]. Cell-type-specific, RPKM-normalized coverage tracks in BigWig format (.bw) were obtained from the WashU Comparative Epigenome Browser[40]. Macaque and marmoset tracks were converted from rheMac10 and calJac4, respectively, to hg38 using the bigwig subcommand of CrossMap (v0.7.0) and the UCSC chainfiles (rheMac10ToHg38.over.chain.gz and calJac4ToHg38.over.chain.gz). No optional arguments were supplied; in the inspected installation, this corresponded to --chromid a, which preserved the input chromosome-ID style. Mapped intervals were written to intermediate bedGraph files, sorted by genomic coordinates, and merged to resolve overlapping entries before conversion to BigWig format. No additional signal normalization was performed. All tracks were deposited in the study's public track hub (https://ccg2025.s3.us-east-2.amazonaws.com/hub.txt), and excitatory-neuron subclasses were visualized in the UCSC Genome Browser[2] (https://genome.ucsc.edu/s/clee03/CCG2026_Fig1B_Exp; **Figure 1B**) to compare promoter-proximal coverage at the disease-associated alternative *RBFOX1* promoter.

**Phylogenetic tree and cross-species genome-wide alignment**

Phylogenetic and comparative genomic analyses used 14 Primatomorpha genome assemblies retrieved from NCBI RefSeq, comprising 13 primates and the Philippine flying lemur (*Cynocephalus volans*) as a dermopteran outgroup (**Table S2**). The human reference was GRCh38.p14 (GCF_000001405.40). A guide phylogeny was inferred using ROADIES[33] (Git commit 6447610) in accurate mode (--mode accurate), with the default fragment length of 500 bp and an initial target of 250 loci (GENE_COUNT=250). The default convergence procedure terminated after the first iteration, yielding 224 gene trees; all evaluated internal branches met the local posterior probability threshold of 0.95. The inferred tree was rooted on *C. volans* using MEGA10[47], and the resulting topology and branch lengths guided the genome-wide alignment. The guide tree was specified as follows:

*(((((((Homo_sapiens:0.00416900,(Pan_troglodytes:0.00057600,Pan_paniscus:0.00052300):0.00299900):0.00204400,Gorilla_gorilla:0.00572500):0.00871200,(Pongo_abelii:0.00242200,Pongo_pygmaeus:0.00085700):0.00340400):0.00241800,Symphalangus_syndactylus:0.02326400):0.01144700,(Macaca_fascicularis:0.00379100,Macaca_nemestrina:0.00439300):0.03438400):0.01572900,(Callithrix_jacchus:0.02331500,Saimiri_boliviensis:0.02065100):0.03646500):0.07642300,(Nycticebus_coucang:0.11490100,Lemur_catta:0.05613400):0.06100900):0.06444200,Cynocephalus_volans:0.06444200);*

Before constructing the genome-wide alignment, primary chromosomal scaffolds in the NCBI assembly reports and associated with valid RefSeq accessions were retained and assigned UCSC-style chromosome identifiers. Alternative loci, assembly patches, and unlocalized and unplaced scaffolds were excluded. The cross-species genome alignment was generated using Progressive Cactus[34] (v3.2.0: Git commit 632f051), managed by Toil (v9.3.0), with local binaries (*--binariesMode local*) and default alignment parameters, including RED repeat masking and CPU-based LASTZ alignment. The resulting alignment was stored in Hierarchical Alignment (HAL) format[48]. For human-reference analyses and visualization, the human genome label was changed from Homo_sapiens to hg38, and the HAL alignment was exported using cactus-hal2maf v3.2.0 with *--refGenome hg38 --outType single --chunkSize 500000 --noAncestors --filterGapCausingDupes*. The exported MAF was converted to bigMaf format using cactus-maf2bigmaf (v3.2.0) for visualization in the UCSC Genome Browser[2].

To assess the consistency of the *RBFOX1* CCG-repeat configuration across alignment resources, we inspected publicly available whole-genome alignments: the 8-way T2T primate CACTUS alignment[3], 241-way and 447-way Zoonomia mammalian CACTUS alignments[49,50], and UCSC MULTIZ alignments of 30 mammals, 470 mammals, and 100 vertebrates[2]. The corresponding *RBFOX1* CCG repeat motif and flanking sequences were summarized as sequence logos (**Figure S1A**; see "Sequence-logo summaries" below).

#### ReAlignPro workflow and local promoter alignments

**ReAlignPro workflow.** ReAlignPro is a Python command-line framework for recovering orthologous promoter intervals, producing local multiple-sequence alignments, identifying species- or group-specific sequence states, and generating transcription start site (TSS)-anchored sequence maps. The framework comprises three subcommands, *fa2maf*, *maf2bed*, and *tsv2fig* (**Figure S2A**), and relies on LASTZ[26] (v1.04.52), MUSCLE[27] (v3.8.1551), and samtools[28] (v1.21). Analyses were performed using ReAlignPro v0.1.1 with Python v3.9.5. Source code is available at https://github.com/chulbioinfo/ReAlignPro.

**Ortholog search and selection (fa2maf).** The genome search set comprised 153 mammalian assemblies from the VGP Phase I v1.0 data freeze[1], including the human hs1 T2T-CHM13 assembly (**Table S2**). Assemblies were retrieved using NCBI Datasets[29] (v16.41.0) with the command *datasets download genome accession <accession> --include genome --no-progressbar*. Human promoter queries were obtained from hg38 and corresponded to chr16:6,018,803-6,019,902 for *RBFOX1* and chrX:23,334,124-23,335,223 for *PTCHD1* (1-based, inclusive coordinates; 1,100 bp each). These search queries were distinct from the shorter TSS-anchored windows used for sequence visualization.

Both promoter queries were searched against the genome set using LASTZ (v1.04.52)[26] with a 12-of-19 spaced seed allowing up to one transition at the designated match positions (*T=1*) and both query strands searched (*--strand=both*). For each assembly, the best-scoring hit was retained as a candidate if it had ≥ 60% sequence identity, covered ≥60% of the query, and had a best-to-second-best hit score ratio of ≥ 1.2. Candidate intervals were extracted using samtools faidx (v1.21), oriented to match the query strand, and aligned back to GRCh38. For every accepted candidate, the best-scoring GRCh38 hit mapped to the query chromosome and overlapped the query interval by ≥ 100 bp and ≥ 20% of its length.

An additional synteny check was applied to *RBFOX1*. The terminal 200-bp sequences at both ends of the query were aligned to the target genome. Both anchors were required to map uniquely, with a best-to-second-best hit score ratio of ≥ 1.2, to the primary-hit contig within 200 kb of the primary-hit midpoint. Candidates that failed only the primary-hit score-ratio criterion could be retained if they passed this synteny check; however, no candidate was retained under this exception.

Orthologous promoter intervals were recovered from 144 of 153 assemblies for *RBFOX1* and 150 of 153 assemblies for *PTCHD1*, including hs1 in both cases. For *RBFOX1*, intervals from *Suncus etruscus, Sorex araneus,* and *Monodelphis domestica* were extended 100-150 bp upstream of the best-hit boundary and re-extracted from the corresponding genomes. Addition of the hg38 query yielded final alignment datasets representing 145 assemblies for *RBFOX1* and 151 for *PTCHD1*, each including both human reference assemblies. Assembly accessions and orthologous interval coordinates are provided in **Table S2**.

**Alignment and TSS anchoring.** Selected intervals and the human query were aligned using MUSCLE[27] (v3.8.1551) with -maxiters 2 -diags, and alignment rows were ordered according to the species order in a mammalian family tree which was constructed in this study using ROADIES[33] (commit 6447610) with the same default options like the 14-way primatomorpha tree construction. The cladogram tree was specified as follows with VGP lineage IDs and academic names:

*(((((((((((((m11_Homo_sapiens,(m11_Pan_troglodytes,m11_Pan_paniscus)),m11_Gorilla_gorilla),(m11_Pongo_abelii,m11_Pongo_pygmaeus)),m11_Symphalangus_syndactylus),(m11_Macaca_fascicularis,m11_Macaca_nemestrina)),m12_Callithrix_jacchus),(m14_Nycticebus_coucang,m15_Lemur_catta)),m16_Cynocephalus_volans),((((((((((m22_Mus_musculus,m22_Apodemus_sylvaticus),m22_Arvicanthis_niloticus),(m22_Micromys_minutus,m22_Rattus_norvegicus)),(m22_Acomys_minous,m22_Acomys_russatus)),(((m59_Chionomys_nivalis,m59_Microtus_pennsylvanicus),m59_Arvicola_amphibius),m59_Onychomys_torridus)),m23_Jaculus_jaculus),(m26_Thomomys_bottae,m27_Castor_canadensis)),(m17_Erethizon_dorsatum,(m19_Muscardinus_avellanarius,((m20_Sciurus_vulgaris,m20_Sciurus_carolinensis),(m20_Marmota_flaviventris,(m20_Urocitellus_parryii,m20_Spermophilus_citellus)))))),(m29_Ochotona_princeps,m28_Oryctolagus_cuniculus)),m30_Tupaia_tana)),(((((((((((((((((((m32_Tursiops_truncatus,m32_Stenella_coeruleoalba),m32_Delphinus_delphis),(m32_Grampus_griseus,(m32_Globicephala_melas,m32_Pseudorca_crassidens))),m32_Orcinus_orca),m32_Lagenorhynchus_albirostris),m32_Lagenorhynchus_acutus),(m32_Monodon_monocero,(m32_Phocoena_sinus,m32_Phocoena_phocoena))),m32_Inia_geoffrensis),(m32_Hyperoodon_ampullatus,(m32_Mesoplodon_densirostris,(m32_Mesoplodon_bidens,m32_Mesoplodon_mirus)))),m32_Kogia_breviceps),(m32_Eubalaena_glacialis,(m32_Balaenoptera_acutorostrata,(m33_Eschrichtius_robustus,(m32_Megaptera_novaeangliae,(m32_Balaenoptera_musculus,m32_Balaenoptera_ricei)))))),m33_Hippopotamus_amphibius),(((m34_Dama_dama,m34_Cervus_elaphus),m34_Muntiacus_reevesi),m34_Bos_taurus)),m35_Sus*

_scrofa),m36_Camelus_dromedarius),(((((((((m42_Mustela_nivalis_vulgaris,m42_Mustela_lutreola),m42_Mustela_erminea),m42_Neogale_vison),m42_Lutra_lutra),m42_Martes_martes),m42_Meles_meles),(m42_Zalophus_californianus,m42_Halichoerus_grypus)),((((m42_Canis_lupus_baileyi,m42_Canis_lupus_orion),m42_Canis_lupus_familiaris),m42_Lycaon_pictus),m42_Vulpes_vulpes)),((m43_Panthera_onca,m43_Neofelis_nebulosa),m43_Lynx_canadensis)),m44_Manis_pentadactyla)),(m37_Equus_caballus,(m38_Diceros_bicornis_minor,m39_Tapirus_indicus))),((((((((((((m40_Pipistrellus_pygmaeus,m40_Pipistrellus_hanaki),m40_Pipistrellus_pipistrellus),m40_Pipistrellus_kuhlii),m40_Pipistrellus_nathusii),m40_Nyctalus_leisleri),m40_Vespertilio_murinus),(m40_Eptesicus_fuscus,m40_Eptesicus_nilssonii)),((m40_Plecotus_auritus,m40_Corynorhinus_townsendii),m40_Antrozous_pallidus)),((((m40_Myotis_myotis,m40_Myotis_nattereri),m40_Myotis_daubentonii),m40_Myotis_emarginatus),m40_Myotis_mystacinus)),m40_Miniopterus_schreibersii),(((m40_Molossus_nigricans,m40_Molossus_molossus),m40_Molossus_alvarezi),(m40_Tadarida_brasiliensis,m40_Mops_condylurus))),(((((m40_Artibeus_lituratus,m40_Artibeus_intermedius),m40_Glossophaga_mutica),m40_Phyllostomus_discolor),m40_Desmodus_rotundus),(m40_Rhynchonycteris_naso,(m40_Saccopteryx_bilineata,m40_Saccopteryx_leptura)))),(((((((m41_Rhinolophus_perniger_lanosus,m41_Rhinolophus_trifoliatus),m41_Rhinolophus_yonghoiseni),m41_Rhinolophus_affinis),(m41_Rhinolophus_hipposideros,m41_Rhinolophus_ferrumequinum)),((m41_Hipposideros_larvatus,m40_Aselliscus_stoliczkanus),m40_Doryrhina_cyclops)),(m41_Rhinopoma_microphyllum,m41_Megaderma_spasma)),m41_Rousettus_aegyptiacus))),(((m45_Sorex_araneus,m45_Suncus_etruscus),m47_Erinaceus_europaeus),m46_Talpa_europaea))),(((((m1_Loxodonta_africana,m1_Elephas_maximus_indicus),m2_Heterohyrax_brucei),(m3_Trichechus_inunguis,m3_Dugong_dugon)),m4_Rhynchocyon_petersi),((m8_Tamandua_tetradactyla,m9_Choloepus_didactylus),m10_Dasypus_novemcinctus))),(((((m49_Trichosurus_vulpecula,m50_

*Macropus_eugenii),m51_Phascolarctos_cinereus),m52_Macrotis_lagotis),m55_Dromiciops_gliroides),m56_Monodelphis_domestica)),(m58_Tachyglossus_aculeatus,m58_Ornithorhynchus_anatinus));*

Conventional gapped alignments and gapless TSS-anchored sequence maps were retained as distinct outputs. Gapped alignments preserve homologous alignment columns, whereas TSS-anchored maps remove alignment gaps and position each sequence relative to the site aligned to the human reference TSS: hg38 chr16:6,019,024 for *RBFOX1* and chrX:23,334,849 for *PTCHD1*. TSS-anchored windows extended 150 bp on either side of this position for *RBFOX1* and 160 bp on either side for *PTCHD1* (**Figure S2B**). Portions of these windows extending beyond the extracted intervals were padded with N. TSS anchoring provides a common positional reference for visualization; it does not establish an experimentally measured TSS in every non-human species. Alignment columns in which the human base was shared by ≥ 95% of the other sequences were flagged as conserved. Gapped alignments were exported in FASTA and MAF formats, whereas TSS-anchored sequence maps were exported in FASTA, MAF, and TSV formats (**Figure S2B**). In the MAF outputs, start coordinates for minus-strand rows were reported in the forward-strand coordinate system.

**Target-specific sites (maf2bed).** The *maf2bed* command identifies reference-coordinate sites that meet target-versus-background sequence-state criteria, using the same general state-set comparison described for ConVarFinder[51]. Let $A_i$ denote the observed state set in the specified target assemblies and $C_i$ the state set in the non-outgroup background assemblies at an evaluable reference position i. An evaluable position was defined as an alignment column in which the hg38 sequence carried a base. Alignment gaps were treated as a state. A site was classified as target-specific when:

$$|A_i| = 1, \quad A_i \cap C_i = \emptyset \qquad (1)$$

All specified target assemblies were required to be present, and consecutive qualifying reference positions were merged into BED intervals. For the human-lineage screen, hg38 and hs1 (T2T-CHM13v2.0; GCA_009914755.4) were specified as the target assemblies. No outgroups were specified; all remaining assemblies therefore formed the background.

**Visualization (tsv2fig).** The *tsv2fig* command renders TSS-anchored TSV matrices as vector PDFs. Variant mode with CCG highlighting was used to distinguish repeat motifs, and homologous regions were manually highlighted (**Figures 1D**, **4A**, and **S2B**). Full ortholog-selection, local-alignment, and TSS-anchored sequence outputs are provided in **Table S2.**

**Sequence-logo summaries**

The homologous sequences of the *RBFOX1* CCG repeat motif (4xCCG.c.CCG) and ±3bp flanking regions (hg38: tgtCCGCCGCCGCCGcCCGctg) were manually extracted as FASTA format from the gapped ReAlignPro local alignment and the independent whole-genome alignment resources described above. Sequence logos were generated using Logomaker[35] (v0.8.7) and custom Python scripts available in the ReAlignPro repository (https://github.com/chulbioinfo/ReAlignPro/). Logos from the independent alignment resources are presented in **Figure S1A**, whereas the ReAlignPro mammalian alignment was summarized separately by lineage in **Figure S1B**, with group sizes indicated. Letter heights represent nucleotide counts at each alignment column, rather than information content.

#### Archaic hominin and modern human variation

ArcSeqHub variant catalogs and AADR[13] (v66.p1) SNP-marker panels were examined at the *RBFOX1* 4xCCG.c.CCG motif (CCGCCGCCGCCGcCCG). The sequence-verified motif interval was hg38 chr16:6,018,926–6,018,941, corresponding to hg19/b37 chr16:6,068,927–6,068,942 (1-based, inclusive coordinates). UCSC-distributed ArcSeqHub Neanderthal and Denisovan hg38 VCFs[12] were queried across the motif to inspect cataloged variation (**Figure 1C**). Marker coordinates from the AADR 1240K, 2M, Human Origins, and compatibility Human Origins *.snp* files were intersected with the motif and its 10-kb flanking regions to assess panel coverage (**Figure S3A**). The human 4xCCG.c.CCG motif was not inferred from the absence of an AADR panel marker or an ArcSeqHub variant record.

Next, read-level support was assessed for the complete 4xCCG.c.CCG motif using published high-coverage archaic genome alignments. BAM files for three Neanderthals (Altai Denisova 5[14], Vindija 33.19[15], and Chagyrskaya 8[16]) and the Denisovan individual Denisova 3[17] were obtained from the Max Planck Institute for Evolutionary Anthropology (**Key resources table** in this **STAR Methods**). Existing alignments were analyzed directly, without FASTQ preprocessing or read remapping. BAM file integrity and indexed access were checked using SAMtools/HTSlib v1.11. Local reference compatibility was verified by comparing the motif and its 50-bp flanking regions across b37, hs37d5, and GRCh38.p14.

Read-level evidence was evaluated using custom scripts written in Python v3.12.2 with pysam (v0.22.1). Primary alignments with mapping quality $\geq 30$ were retained, excluding MAPQ 255. Unmapped, secondary, supplementary, QC-failed, and duplicate-marked records were excluded using the SAM flag exclusion mask 0xF04. Complete-repeat assessment required each read to span the intervening allele and two contiguous 10-bp anchors immediately flanking it, with both anchors matching the reference exactly. Every base in the anchors and intervening allele was required to have a Phred quality score ≥ 20. Allele sequences were extracted using CIGAR-derived reference-to-query mappings while preserving observed insertions and deletions. Reads with incomplete anchors, reference skips, empty alleles, or non-ACGT bases in the allele were excluded from complete-repeat scoring.

Read observations sharing library-qualified read identifiers or inferred fragment endpoints were grouped into putative molecule groups. Read-group identifiers were used when library identifiers were unavailable. Observed alleles were classified as the exact 16-bp reference sequence, the exact 13-bp contracted sequence (CCGCCGCCGcCCG), another 16-bp sequence containing substitutions, or another length. Structural support required at least three putative molecule groups supporting the exact 16-bp reference sequence, with no fully anchored alternative-length observations and no within-group conflicts. Insufficient evidence was designated NO CALL. Observed substitutions were retained without automatic correction for ancient-DNA damage, and each individual was evaluated separately.

**Figures S3B–D** were generated using Matplotlib v3.10.1. Quality-filtered per-base depth and all motif-overlapping alignments retained after the alignment-level filters described above were displayed without downsampling (**Figure S3B**). Complete-repeat support and observed allele sequences were summarized separately after grouping read observations into putative molecule groups (**Figures S3C** and **S3D**, respectively). In these datasets, each passing anchored read belonged to a distinct putative molecule group. These counts summarized sequence evidence under the specified filters and were not interpreted as diploid genotypes or population allele frequencies.

**Transcription-factor motif scanning**

Predicted transcription-factor binding sites in the *RBFOX1* promoter were first inspected using the precomputed JASPAR CORE 2022 track in the UCSC Genome Browser (**Figure 2A**). These genome-wide predictions had been generated using PWMScan. For visualization, the minimum UCSC track score was set to 350, corresponding approximately to $P \leq 3.16 \times 10^{-4}$. Predicted binding-site positions were examined in relation to the CCG-repeat motif and the 14-way Primatomorpha genome alignment.

To crosscheck the browser-based predictions and compare motif occurrences among orthologous *RBFOX1* alleles, sequences spanning the CCG-repeat region were analyzed using FIMO v5.5.9 (MEME Suite) with the 841 matrices from the JASPAR2022 CORE vertebrates non-redundant collection in MEME format. The input comprised 144 ungapped sequences from 143 non-human mammalian assemblies and the human hs1 assembly (**Table S2**). The human input sequence, tgtCCGCCGCCGCCGcCCGctg, comprised the 16-bp 4xCCG.c.CCG motif and 3 bp of flanking sequence on each side, spanning hg38 chr16:6,018,923-6,018,944 (1-based, inclusive coordinates; 22 bp). Because hg38 and hs1 were identical across this interval, the human sequence was included only once. Homologous non-human input sequences are provided in **Table S2**.

Both DNA strands were scanned using *--thresh 1e-4 --max-strand*, with motif occurrences reported at an unadjusted P-value $< 10^{-4}$ When both strands at the same position passed this threshold, only the higher-scoring strand was reported. The analysis used FIMO's default zero-order nucleotide background (A = T = 0.275; C = G = 0.225) and a motif pseudocount of 0.1. FIMO-generated q-values were retained for reporting, with hit inclusion determined by the P-value threshold. *EGR1* and *WT1* predictions were identified using the

MA0162.4 and MA1627.1 matrices (JASPAR label: *Wt1*), respectively. Predicted positions and matched sequences were compared with the UCSC annotations and among orthologous alleles. The complete set of 673 FIMO motif occurrences, including scores, P-values, q-values, and matched sequences, is provided in **Table S3**.

**Genome-wide motif catalog and human-derived candidates**

Genome-wide occurrences of the 16-bp sequence 5'-CCGCCGCCGCCGcCCG-3' were cataloged from a local BLAST+ (v2.16.0+) search of the GRCh38.p14 RefSeq assembly (GCF_000001405.40), retaining full-length exact matches on both strands. The catalog was subsequently validated by exhaustive, case-insensitive searches for the motif and its reverse complement across all sequences in the assembly, including soft-masked regions and overlapping occurrences. These searches recovered the same 266 assembly-coordinate occurrences, comprising 132 forward-strand and 134 reverse-strand matches. Of these, 257 occurred on assembled chromosomes, two on an unlocalized scaffold, and seven on patch scaffolds. Patch-scaffold occurrences were retained as separate assembly-coordinate records.

RefSeq sequence accessions were mapped to UCSC sequence names using the NCBI Assembly Sequence Report. Coordinates were normalized to 0-based, half-open intervals and sorted using BEDTools[31] (v2.31.1). Core-promoter intervals were defined from transcript-level TSSs in the NCBI RefSeq annotation GCF_000001405.40-RS_2025_08. Each interval included the TSS base and 150 bp on either side, yielding a 301-bp window. A custom Python interval-overlap script associated motif loci with core-promoter intervals sharing at least 1 bp, irrespective of strand. Gene symbols were deduplicated separately within each motif locus. This procedure identified 104 core-promoter-associated motif loci, yielding 108 locus-gene associations involving 104 distinct gene symbols (**Figure 3A**; **Table S4**).

To screen for human-derived candidates, the 16-bp motif intervals were compared with human-specific annotation intervals derived from the 14-way Primatomorpha alignment. For each motif, the number of overlapping bases and their fraction of the 16-bp interval were recorded. Motifs overlapping these annotations by at least 1 bp were designated candidate loci, yielding 62 candidates, including 20 core-promoter-associated loci representing 19 distinct gene symbols. Human-chimpanzee motif-region alignments for the core-promoter candidates are provided in **Table S4**. Three *RGPD1*-associated core-promoter loci had entirely gapped chimpanzee alignment entries and were classified as unresolved with respect to alignment and orthology. Locus-level coordinates and gene associations were retained separately for these duplicated-region candidates. These three unresolved loci were excluded from downstream variant-effect analyses, leaving 17 core-promoter candidates.

The chromosome map in **Figure 3A** was generated in R using *ggplot2* and *ggrepel*. The annotated motif BED file and hg38 chromosome-length data were imported using readr and processed using *dplyr*. Plotting was restricted to core-promoter-associated motif intervals on chromosomes 1-22, X, and Y, with each locus represented by a point at the midpoint of its BED interval. Chromosomes were drawn to scale in a paired horizontal layout with a 50-Mb scale bar. For identical core-promoter annotation strings recurring on the same chromosome, a single label was anchored at the median of the corresponding locus midpoints, while all individual loci were retained as points. Labels were positioned using ggrepel with vertical displacement and a fixed random seed of 1. The map included 103 primary-chromosome motif loci. The

*DGKD*-associated fix-patch locus (chr2_KN538363v1_fix:299,904-299,920) was retained in **Table S4** but excluded from the chromosome map.

**Functional enrichment and ASD-associated expression**

Functional enrichment of the 104 unique genes associated with motif-bearing core promoters (**Table S4**) was assessed using g:Profiler g:GOSt, with Homo sapiens selected as the organism. Overrepresentation was evaluated using the default hypergeometric test against the annotated human genes in each data source. P values were adjusted using g:SCS, and significance was defined as adjusted $P < 0.05$. **Figure 3B** shows all significant terms from the GO biological-process (n = 13), GO cellular-component (n = 5), and Human Protein Atlas (n = 1) categories. A custom Python script grouped terms by annotation source and ordered them by decreasing $-\log_{10}$(adjusted P value). Horizontal position indicates $-\log_{10}$(adjusted P value), and bubble size indicates the number of query genes annotated to each term.

The 16 genes with motif-bearing core promoters that were annotated in SFARI Gene (2025 Q4) were selected for ASD-associated expression analysis (**Figure 3C**; **Table S4**). Published cell-subtype-specific ASD-versus-control differential-expression statistics from the PsychENCODE frontal-cortex single-nucleus RNA-sequencing resource were retrieved through the ASD Single-Cell Gene Expression Portal (ASD scGENE Portal)[24]. Records corresponding to these genes were extracted by exact gene-symbol matching. Portal-provided log fold changes and FDR values were used directly; differential-expression models were not refitted, and multiple-testing corrections were not recalculated.

Expression patterns were visualized using a custom R script with *ggplot2*. Bubble fill represented the reported ASD-versus-control log fold change, with negative values shown in blue, zero in white, and positive values in red; the color scale saturated at -2 and +2. Bubble size represented $-\log_{10}$(FDR), capped at 10 for display, and thicker black outlines indicated FDR < 0.05. Both significant and nonsignificant observations were displayed for the 16 genes, with genes ordered alphabetically and cell subtypes grouped by their annotation prefixes.

#### AlphaGENome-Integrated Explorer (Alpha-GENIE)

**AlphaGenome variant definitions and analysis windows.** Variants were represented on the forward strand of GRCh38 using 1-based, VCF-style positions and explicit REF and ALT alleles. For *RBFOX1* and the genome-wide promoter-candidate analysis, REF denoted the human reference allele and ALT the chimpanzee-like allele. For *PTCHD1*, REF denoted the human reference allele and ALT the autism-associated duplication allele. The underlying allelic comparison was therefore consistently oriented as ALT relative to REF; differences in biological interpretation arose from the allele definitions, not from a reversal of the comparison direction.

The *RBFOX1* variant was represented as chr16:6,018,925, REF T**CCG** and ALT T, corresponding to deletion of chr16:6,018,926-6,018,928. **Figures 2B** and **2C** used the 16,384-bp input interval chr16:6,010,734-6,027,118 (0-based, half-open coordinates). **Figures 3D-F** used 1,048,576-bp input intervals for the 17 variants remaining after exclusion of the three *RGPD1*-associated records from the initial 20-candidate set. The *RBFOX1* result from this 1-Mb analysis was analyzed separately from its 16-kb result. **Figures 4B** and **4C** used the *PTCHD1* variant at chrX:23,334,765, with REF T and ALT T**CGCCGCCGCCGCGGGCGCCGCTGCCGC**, representing a net 27-bp insertion, within the 1,048,576-bp input interval chrX:22,810,477-23,859,053. Complete allele definitions and fixed input intervals for the candidate set are retained in the versioned input and matched-null manifests.

**Reference checks and target gene assignment.** REF alleles were checked against the local hg38 FASTA. GENCODE annotations were used to assess chromosome agreement, TSS inclusion in the input interval and distance from the edited interval center, or insertion point, to the intended gene's TSS. The target-assignment procedure prioritizes MANE Select transcripts on the variant chromosome, otherwise the nearest annotated transcript, with a gene-feature fallback when transcript annotations are unavailable. It records membership in a ±250-bp TSS window and a strand-aware proximal-promoter window from -2,000 to +500 bp. The saved QC records identify ENST00000550418.6 for *RBFOX1* and ENST00000379361.5 for *PTCHD1*. These checks document positional plausibility; they do not establish a causal target gene. Neighbor-gene score summaries are QC information and do not replace the specified target. Transcript selection for annotation is distinct from the provider's gene exon mask used for scoring. The local distribution specifies hg38 FASTA and GENCODE v46 GTF inputs and records the configured reference identity.

**AlphaGenome RNA effect scoring.** AlphaGENIE (AlphaGENome-Integrated Explorer) was used for matched-null processing and visualization. Human RNA_SEQ predictions were obtained with AlphaGenome Python SDK 0.8.0 using GeneMaskLFCScorer (requested_output=RNA_SEQ)[25]. Target-gene rows were selected by gene symbol, output type and scorer identity. The returned raw_score is a signed predicted log-fold-change statistic over a gene exon mask, not a difference of untransformed coverage or a measured change in expression. The API client does not expose the logarithm base; scores were therefore retained in the returned units and were not relabeled as log2 fold change or percentage change. Quantile scores were auxiliary annotations. Other prediction modalities were not used in the primary RNA-effect analysis.

**Matched null variant generation.** Each of the 19 variant-by-window contexts was evaluated against 1,000 pseudo-variants, generated with NumPy random seed 20260527 and scored for the same target gene within the real variant's fixed input interval. Null intervals were not recentered on the pseudo-variant positions. The real variant and duplicate candidate representations were excluded; candidate anchors or edited reference sequences containing N were rejected. Sampling and scoring had to produce the requested number of valid nulls.

Pseudo-deletions preserved deletion length. Sampling proceeded through four stages as needed: deleted sequence containing CG with an absolute GC-fraction difference from the real deleted sequence of at most 0.10; CG-containing sequence with tolerance 0.20; tolerance 0.20 without a CG requirement; and length-only matching. GC was calculated over the deleted sequence, not a flanking genomic window. Candidate deletions were left-normalized within the input interval, with a maximum 100-bp left shift, before deduplication. The *RBFOX1* 16-kb analysis used 296 first-stage controls and 704 controls from the stage without a CG requirement. Accordingly, these 1,000 controls were not all CG-matched.

Pseudo-substitutions initially matched the full REF/ALT pattern. If necessary, unedited bases were allowed to vary while preserving the original reference and replacement bases at the edited offsets. Joint multi-nucleotide substitutions were retained as joint edits. *PTCHD1* pseudo-insertions placed the same 27-bp inserted sequence at other valid reference anchors. These controls calibrate sequence-class effects within a fixed genomic context; they do not comprehensively match TSS distance, CpG-island overlap, repeat phase or mappability. Per-null identities, coordinates and sampling stages were retained.

**Trackwise matched-null calibration.** Let r denote the raw GeneMaskLFC score for real variant v and canonical RNA track t, and let the superscript null denote the score for its kth matched pseudo-variant. For N = 1,000 null variants, the trackwise null median b was subtracted from both the real and null scores to obtain adjusted effects a:

$$b_{vt} = median_{k=1,\dots,N}(r^{null}_{vkt})$$

$$a_{vt} = r_{vt} - b_{vt};\ a^{null}_{vkt} = r^{null}_{vkt} - b_{vt} \quad (2)$$

The same trackwise center was applied to every null without leave-one-out recentering. This calibration removes the sampled track-specific background median; it does not establish that all model bias has been removed. Category aggregation and empirical inference are described under Quantification and statistical analysis.

**RNA track classification and completeness.** The versioned Brain9 catalog contains 371 RNA tracks. Canonical track identity combines track name, biosample name, ontology identifier and data source; biosample material, donor life stage and GTEx tissue metadata are also checked. The real variant and every null must contain all 371 reviewed track identities with finite scores. Missing or unknown tracks, changed metadata, nonfinite values, or discrepancies between sampled and scored null IDs stop analysis. Duplicate real rows are collapsed by track median and duplicate null rows by the median within each null-ID and track combination before calibration. No missing-track imputation or substitution from another dataset is performed.

Brain9 comprises eight adult anatomical categories containing 14 tracks and one pooled Embryo category containing seven embryonic brain-tissue tracks. Non-brain tissues and Cells and cell lines are separate comparison pools across available donor stages and are excluded from Brain9 inference and cosine similarity.

| CATEGORY | TRACKS | ROLE |
|---|---|---|
| Cerebral cortex | 4 | Adult Brain9 |
| Hippocampal formation | 1 | Adult Brain9 |
| Basal ganglia and striatum | 3 | Adult Brain9 |
| Amygdala | 1 | Adult Brain9 |
| Diencephalic regions | 1 | Adult Brain9 |
| Midbrain | 1 | Adult Brain9 |
| Cerebellum | 2 | Adult Brain9 |
| Whole brain | 1 | Adult Brain9 |
| Embryo | 7 | Embryonic Brain9 |
| Non-brain tissues | 153 | Display only |
| Cells and cell lines | 197 | Display only |

Embryo pools four cortex, one diencephalon and two cerebellum tracks directly, rather than first averaging these regions. Whole brain denotes a region-unspecified adult brain track, not an average of the other categories. Cultured neural/glial cells, embryonic stem-cell models and neural tumor lines are assigned to Cells and cell lines; spinal cord, pituitary and peripheral nerve are assigned to Non-brain tissues. Track counts denote assay outputs, not independent biological replicates.

**Frontal cortex curves and figure generation.** The locus-level RNA curves use the unique adult frontal cortex track identified by GTEx Brain_Cortex, UBERON:0001870, polyA plus RNA-seq and unstranded orientation. This selection is distinct from Whole brain, BA9 and embryonic cortex. REF and ALT channels are matched independently by metadata and must cover the requested interval at 1-bp resolution. A missing or ambiguous match produces an unavailable curve rather than a substituted tissue; it does not invalidate otherwise complete matched-null scores.

Raw arrays are preserved before reference-coordinate projection. For deletions, only deleted reference positions are set to zero in the projected ALT array. For insertions, the shared anchor and inserted segment are collapsed by their maximum; downstream ALT indices are shifted by the net insertion length, and unmappable boundaries remain missing. These operations support coordinate-aligned visualization and do not alter the gene-level scorer. Gene structures are drawn from the local annotation, prioritizing MANE Select, Ensembl canonical and APPRIS principal transcripts, followed by exon length. The curves display positional REF and ALT predictions and their direct difference; the bars display matched-null-adjusted GeneMaskLFC scores.

The *RBFOX1* curve displays the full 16,384-bp interval and a partial gene model. The *PTCHD1* curve displays an 89,526-bp gene-centered segment, chrX:23,324,848-23,414,374 in 0-based, half-open coordinates, extracted from its 1,048,576-bp prediction interval. Saved curves retain the union of REF, ALT and difference minima/maxima from 1,024 contiguous bins, together with endpoints and edit-adjacent positions, without averaging. The difference axis uses a symmetric-log scale.

Single-variant figures show category medians as bars and canonical tracks as points, with track counts in category labels. **Figure 3** displays the nine Brain9 columns in a heatmap, consensus effects with FDR annotations, and similarity to *RBFOX1*. Default rows are ordered by increasing Brain9 consensus, not genomic position or clustering. The web interface separates gene-structure/curve and category-effect panels vertically and exposes track metadata on hover.

For saved single-variant results, user-configured PDF exports stack these panels. Saved manuscript figure layouts and source arrays are preserved separately from interactive export settings.

**Public and local deployment.** AlphaGENIE v0.24 provides a browser interface for saved results, with publicly available source code in the v0.24.0 distribution (https://github.com/chulbioinfo/alphagenie; https://alpha-genie.net). Viewing or exporting saved results does not request new AlphaGenome inference; public new-analysis submission is disabled. Local users can submit new analyses with their own API key and local reference files. Data processing, storage and visualization occur locally, whereas authorized prediction requests are sent directly to Google DeepMind. AlphaGENIE operators do not collect these credentials. Optional key storage is an owner-only plaintext file outside the repository. The local server is intended for one trusted OS account on a personal computer and has no multi-user authentication or isolation boundary. Use is subject to the applicable AlphaGenome API and output terms; predictions are research outputs, not clinical diagnoses.

**Promoter–luciferase reporter construction**

Matched *RBFOX1* promoter reporters encoded the human five-copy allele (4xCCG.c.CCG), chimpanzee-like four-copy allele (3xCCG.c.CCG), ring-tailed lemur allele (3xCCG.c.2xCCG), or an engineered motif-null allele (**Figures 2D** and **S4A**). Chimpanzee-like and motif-null sequences were obtained as single-stranded IDT oligonucleotides with 5’ gacagctggacgtcgat and 3’ atcgaattcgggtatataat cloning adapters. The pBV-Luc backbone (Addgene, 16539) was digested with EcoRV overnight. Promoter oligonucleotide (5 ng) and digested vector (50 ng) were assembled in a 10-μL NEB HiFi reaction; 1 μL was used to transform 25 μL of competent DH5α

E. coli. Individual colonies were miniprepped (Qiagen), screened by EcoRV/XbaI digestion, and sequence-verified using Plasmid-EZ (Genewiz). Because repeated cloning attempts yielded deletions in the GC-rich human tract, the human reporter was obtained by Thermo Fisher GeneArt Complex Gene Synthesis and then transformed and miniprepped.

*PTCHD1* promoter reporters were obtained through the same GeneArt service and encoded the reference allele, the ASD-associated 27-bp duplication allele, or a motif-null allele lacking the 4xCCG.c.CCG motif sequence (**Figures 4D** and **S4C**). The experiment compares the full patient-associated duplication with REF; it does not selectively isolate the added 5xCCG motif from the remaining duplicated bases.

**Transcription-factor expression constructs**

pcDNA3-*EGR1* was obtained from Addgene (11729). pcDNA3-WT1 and pcDNA3-GFP were generated by EcoRI/ApaI digestion of pcDNA3-EGR1, gel purification of the backbone, and HiFi assembly with IDT gBlocks containing the corresponding coding sequence, with 5′ tgtgctggaattctgcagatatctt and 3′ agggccctattctatagtgtcaccta adapters. Assembly products were transformed, miniprepped, and sequence-verified. DNA-binding-domain-deletion constructs (*EGR1ΔBD* and *WT1ΔBD*) were generated by Q5 PCR and HiFi assembly. The existing record identifies deleted amino-acid intervals C349–H427 in *EGR1* and P184–H291 in *WT1*, based on NCBI Conserved Domain Search.

The recorded deletion primers (5′–3′) were *EGR1* forward, TTAAGACAGAAGGACAAGAAAGCAG; *EGR1* reverse, CTGCTTTCTTGTCCTTCTGTCTTAAAGCATATGGGCGTTCATGGG; *WT1* forward,

CAGAGAAACATGACCAAACTCCAG; and *WT1* reverse, CTGGAGTTTGGTCATGTTTCTCTGGTAAGCACACATGAAGGGGC.

**Luciferase reporter assays**

Plasmid concentrations were adjusted to 100 ± 5 ng/µL using a NanoDrop One spectrophotometer. Reporter plasmids and transcription factor expression plasmids were mixed at a 5:2 mass ratio. For each well, 200 ng total plasmid DNA was diluted in 25 µL Opti-MEM (Gibco, 31985070) and combined with 25 µL of an Opti-MEM dilution of Lipofectamine 2000. Three separate 50-µL complexing reactions were prepared for each reporter-by-expression condition. Complexes were formed for at least 20 min.

Cells were suspended in Opti-MEM and counted with acridine orange/propidium iodide (Logos Biosystems, F23001) using a LUNA BX7. Each well received 100 µL of a $7.5 \times 10^5$ live-cells/mL suspension (75,000 cells) and was incubated for 30–36 h at 37°C and 5% CO2. ONE-Glo reagent (150 µL; Promega, E6110) was added, plates were agitated for 5 min at room temperature, and chemiluminescence and brightfield images were acquired using an Amersham 6000 Imager. Mean luminescence per well was measured with custom FIJI/ImageJ macros.

*RBFOX1* and *PTCHD1* reporters were tested with CMV-driven *EGR1*, *EGR1ΔBD*, *WT1*, *WT1ΔBD*, or *GFP*, and with a water-only condition lacking an added expression plasmid. Representative layouts and complete condition matrices are shown in **Figures S4A** and **S4B** for *RBFOX1* and **Figures S4C** and **S4D** for *PTCHD1*. Three technical wells per condition were averaged within each independent transfection experiment; technical wells were not treated as independent experimental replicates. The current **Figure 2E** reports five independent *RBFOX1* transfections.

### QUANTIFICATION AND STATISTICAL ANALYSIS

#### Reporter normalization and independent experimental units

Reporter data were processed at the plate/independent-transfection level according to the current **Figure S4** legend. For plate p, let $B_p$ be the mean signal in no-reporter wells, $L_{part}$ the measured signal for reporter allele a, expression condition r, and technical well t, and $T_{par}$ the number of technical wells. The background-subtracted mean and motif-null+GFP-normalized log2 activity were defined as:

$$\bar{L}_{par} = \frac{1}{T_{par}} \sum_{t=1}^{T_{par}} \left( L_{part} - B_p \right) \qquad (3)$$

$$x_{par} = log_2\left( \frac{\bar{L}_{par}}{\bar{L}_{p,null,GFP}} \right) \qquad (4)$$

Here “null” denotes the engineered motif-null reporter. Background subtraction preceded normalization, and technical replicates were averaged before log2 transformation. Each plotted point represents one independent transfection. Mean log2 activity and its 95% confidence interval were shown when at least three independent experiments were available. Conditions with fewer than three experiments were descriptive (**Figures S4B** and **S4D**).

#### Paired allele contrasts and *EGR1* interactions

Allelic differences under *EGR1* were calculated within matched plates. For an allele pair *a* and *b*, the paired log2 ratio and back-transformed geometric mean fold change were:

$$q_p^{a:b} = x_{pa,EGR1} - x_{pb,EGR1}, \quad FC_{a:b} = 2^{\frac{1}{n}\sum_{p=1}^{n} q_p^{a:b}} \quad (5)$$

Here n is the number of independent plates with a complete pair. Two-sided paired t tests on log2 activity, equivalently one-sample t tests of the paired log2 ratios against zero, were used. The three specified *EGR1* contrasts formed a separate Holm-adjustment family for each gene: chimpanzee-like versus motif-null, human versus chimpanzee-like, and human versus motif-null for *RBFOX1*; REF versus motif-null, ASD versus REF, and ASD versus motif-null for *PTCHD1* (**Figures 2F** and **4F**).

For mean paired log2 ratio $\bar{q}$ and sample standard deviation $s_q$, a t-based 95% interval on the fold-change scale is shown below:

$$95\%\ CI = \left[ 2^{\bar{q} - t_{0.975,n-1}\frac{s_q}{\sqrt{n}}},\ 2^{\bar{q} + t_{0.975,n-1}\frac{s_q}{\sqrt{n}}} \right] \quad (6)$$

*EGR1* induction was calculated within each allele and plate relative to its *GFP* control. The allele-by-*EGR1* interaction was the difference between the two induction values:

$$I_{pa} = x_{pa,EGR1} - x_{pa,GFP} \quad (7)$$

$$J_p^{a:b} = I_{pa} - I_{pb}, \quad FC_{interaction} = 2^{\frac{1}{n}\sum_{p=1}^{n} J_p^{a:b}} \quad (8)$$

For *RBFOX1*, a and b denote human and chimpanzee-like alleles; for *PTCHD1*, a and b denote ASD and REF alleles. The interaction was tested against zero using a separate two-sided paired t test and was not included in the three-contrast *EGR1* Holm family (**Figures 2G** and **4G**). The back-transformed mean interaction is the geometric mean ratio of *EGR1/GFP* induction ratios.

##### AlphaGenome group summaries and empirical tests

For each variant v, the category effect G was the median of adjusted track scores within category g. The consensus C was the median of the nine Brain9 category effects:

$$G_{vg} = median_{t\in g}(a_{vt})\ ;\ \ C_v = median_{g\in Brain9}(G_{vg}) \quad (9)$$

Identical aggregation was applied to each matched null after the trackwise calibration in Equation (2). Each category contributes one value irrespective of its track count; the consensus is not the median of all 21 brain tracks. Non-brain tissues and Cells and cell lines are excluded. RNA tracks are correlated model outputs; the empirical reference units are matched pseudo-variants, not biological samples.

For each variant, m denotes the median of its N = 1,000 null-consensus values. The primary two-sided empirical P value compares the absolute deviations of the real and null consensuses from this median:

$$P_v^{two} = \frac{1 + \sum_{k=1}^{N} I\{|C_{vk}^{null} - m_v| \geq |C_v - m_v|\}}{N+1} \quad (10)$$

The lower- and upper-tail empirical P values were calculated as follows:

$$P_v^{decrease} = \frac{1 + \sum_{k=1}^{N} I\{C_{vk}^{null} \leq C_v\}}{N+1} \quad (11)$$

$$P_v^{increase} = \frac{1 + \sum_{k=1}^{N} I\{C_{vk}^{null} \geq C_v\}}{N+1} \quad (12)$$

The indicator I equals one when its condition is satisfied and zero otherwise. The plus-one correction gives a minimum attainable P value of 1/1,001. The observed-direction tail probability uses the upper tail when the real consensus minus the null median is nonnegative and the lower tail otherwise. Because the tail is selected after observing the effect, these directional values are exploratory rather than prespecified one-sided evidence. The displayed effect is C; subtraction of m is specific to testing and the robust score.

For **Figure 3**, Benjamini–Hochberg adjustment was applied across the complete 17-variant family, separately for the primary two-sided P values and the exploratory observed-direction values. Variants with either adjusted value ≤ 0.05 were labeled in the middle panel, with the primary two-sided and exploratory observed-direction statistics distinguished. Standalone *RBFOX1* and *PTCHD1* analyses report empirical P values without the 17-variant correction. New multi-variant jobs produce cohort inference only when every requested variant succeeds; failures do not reduce the testing family.

A descriptive robust calibration score was calculated from the median absolute deviation (MAD) of the null consensus:

$$z_v = \frac{C_v - m_v}{1.4826 \times MAD(C_{vk}^{null})} \quad (13)$$

The robust score is missing when the null MAD is zero and is not substituted for empirical P values. Missing or nonfinite required track scores, inconsistent null identities, or fewer than the requested number of complete nulls stop analysis.

#### Similarity to the *RBFOX1* brain effect profile

For **Figure 3**, cosine similarity was calculated from the nine uncentered category effects, using RBFOX1_del3 from the 1,048,576-bp cohort as reference R:

$$cos(v,\ R) \ = \ \frac{\sum_{g=1}^{9} G_{vg}\, G_{Rg}}{\sqrt{\sum_{g=1}^{9} G_{vg}^{\ 2}}\sqrt{\sum_{g=1}^{9} G_{Rg}^{\ 2}}} \quad (14)$$

All nine category values must be finite, and categories are not dropped pairwise. A zero-norm vector yields NA rather than zero. Non-brain tissues and Cells and cell lines do not contribute. General local comparisons use the first input row as the reference; reproduction of **Figure 3** therefore requires its *RBFOX1* reference. Cosine similarity describes modeled effect profiles and is not an independent test of shared regulation or biological coexpression.

Candidate prioritization combined a negative consensus brain effect with exploratory observed-direction FDR < 0.05, positive cosine similarity to *RBFOX1* (> 0), and SFARI annotation, applying the same rule to all 17 loci. This prioritization does not replace primary two-sided screen-level inference.

**Other statistical reporting and reproducibility**

Functional enrichment and the PsychENCODE analysis used their analysis-specific multiple-testing procedures (**Figure 3B** and **3C**). AlphaGENIE variant definitions, real/null summaries, track assignments, group statistics, testing families and software/environment records are documented in the versioned release specified under Data and code availability and Computational provenance and verification.

#### ADDITIONAL RESOURCES

Interactive genomic context is available through the study UCSC hub (https://ccg2025.s3.us-east-2.amazonaws.com/hub.txt) and the figure-specific UCSC sessions cited in the legends. ReAlignPro (https://github.com/chulbioinfo/ReAlignPro) and AlphaGENIE (https://github.com/chulbioinfo/alphagenie; https://alpha-genie.net) provide the analysis and visualization entry points.

#### Methods reference

7. Cui, X., Yang, H., Cai, C., Beaman, C., Yang, X., Liu, H., Ren, X., Amador, Z., Jones, I.R.,

Keough, K.C., et al. (2025). Comparative characterization of human accelerated regions in neurons. Nature *640*, 991–999.

13. Mallick, S., Micco, A., Mah, M., Ringbauer, H., Lazaridis, I., Olalde, I., Patterson, N., and Reich, D. (2024). The Allen Ancient DNA Resource (AADR) a curated compendium of ancient human genomes. Sci. Data *11*, 182.

26. Harris, R.S. (2007). Improved pairwise alignment of genomic DNA.

32. Ye, J., McGinnis, S., and Madden, T.L. (2006). BLAST: improvements for better sequence analysis. Nucleic Acids Res. *34*, W6–W9.

33. Gupta, A., Mirarab, S., and Turakhia, Y. (2025). Accurate, scalable, and fully automated inference of species trees from raw genome assemblies using ROADIES. Proc. Natl. Acad. Sci. U. S. A. *122*, e2500553122.

34. Armstrong, J., Hickey, G., Diekhans, M., Fiddes, I.T., Novak, A.M., Deran, A., Fang, Q., Xie, D., Feng, S., Stiller, J., et al. (2020). Progressive Cactus is a multiple-genome aligner for the thousand-genome era. Nature *587*, 246–251.

35. Tareen, A., and Kinney, J.B. (2020). Logomaker: beautiful sequence logos in Python. Bioinformatics *36*, 2272–2274.

36. Grant, C.E., Bailey, T.L., and Noble, W.S. (2011). FIMO: scanning for occurrences of a given motif. Bioinformatics *27*, 1017–1018.

37. Reimand, J., Kull, M., Peterson, H., Hansen, J., and Vilo, J. (2007). g:Profiler--a web-based toolset for functional profiling of gene lists from large-scale experiments. Nucleic Acids Res *35*, W193–W200.

47. Kumar, S., Stecher, G., Li, M., Knyaz, C., and Tamura, K. (2018). MEGA X: molecular evolutionary genetics analysis across computing platforms. Mol. Biol. Evol. *35*, 1547.

48. Hickey, G., Paten, B., Earl, D., Zerbino, D., and Haussler, D. (2013). HAL: a hierarchical format for storing and analyzing multiple genome alignments. Bioinformatics *29*, 1341–1342.

49. Zoonomia Consortium (2020). A comparative genomics multitool for scientific discovery and conservation. Nature *587*, 240–245.

50. Kuderna, L.F.K., Ulirsch, J.C., Rashid, S., Ameen, M., Sundaram, L., Hickey, G., Cox, A.J., Gao, H., Kumar, A., Aguet, F., et al. (2024). Identification of constrained sequence elements across 239 primate genomes. Nature *625*, 735–742.

51. Lee, C., Cho, S., Kim, K.-W., Yoo, D., Davenport, M., Han, J.Y., Lee, H.J., Gedman, G., Audet, J.-N., Hara, E., et al. (2025). Insights into Convergent Evolution From Studying Amino Acid Patterns in Independent Lineages of Birds. Genome Biol Evol *17*, evaf112.
